# WindSTORM: Robust online image processing for high-throughput nanoscopy

**DOI:** 10.1101/434415

**Authors:** Hongqiang Ma, Jianquan Xu, Yang Liu

## Abstract

High-throughput nanoscopy becomes increasingly important for unraveling complex biological processes from a large heterogeneous cell population at a nanoscale resolution. High-density emitter localization combined with a large field of view and fast imaging frame rate is commonly used to achieve a high imaging throughput, but the image processing speed in the dense emitter scenario remains a bottleneck. Here we present a simple non-iterative approach, referred to as WindSTORM, to achieve high-speed high-density emitter localization with robust performance for various image characteristics. We demonstrate that WindSTORM improves the computation speed by two orders of magnitude on CPU and three orders of magnitude upon GPU acceleration to realize online image processing, without compromising localization accuracy. Further, due to the embedded background correction, WindSTORM is highly robust in the presence of high and non-uniform background. WindSTORM paves the way for next generation of high-throughput nanoscopy.

## Introduction

Super-resolution localization microscopy, such as stochastic optical reconstruction microscopy [STORM] and (fluorescence) photo-activated localization microscopy [(f)PALM] ^1–4^, has become an important tool to visualize molecular structures at a nanoscale resolution. Despite its superior spatial resolution, it requires precise localization of sparsely excited single molecules from thousands of frames, which significantly compromises temporal resolution and imaging throughput. High-density emitter localization is an effective strategy to improve the throughput by increasing the emitter density at each frame with reduced number of imaging frames. To precisely localize the overlapping molecules in dense emitter scenarios while maintaining a high localization accuracy^5^, complex numerical optimization algorithms are required^6–12^. As they are computationally intensive, the long image processing time limits their usage to mostly small image size and short temporal sequence. But the heterogeneous nature of cell population often requires high-throughput super-resolution imaging on a large number of cells^13^. Recent advance in sCMOS camera technology has greatly benefited super-resolution microscopy with a large field of view^14–16^ and fast frame rate^6^, and next-generation high-throughput nanoscopy has become feasible. But as high-throughput nanoscopy can routinely generate gigabytes of dataset in seconds, real-time image processing becomes a major challenge. The slow speed of current high-density image processing methods can no longer meet the increasing demand for high-throughput analysis of a huge dataset, while those high-speed image processing methods for sparse emitter scenarios fail in accuracy with unacceptable image quality for high-density data^7,11,17^. Further, in many high-density scenarios, heterogeneous background is present and can induce significant image artifacts. Therefore, an online high-density image processing method that is fast and robust to reconstruct high-quality super-resolution image is crucial for next-generation high-throughput nanoscopy.

We present a computationally simple solution for high-density emitter localization, referred to as WindSTORM, to enable online image processing essential for high-throughput nanoscopy and remain robust even for heterogeneous background. Unlike conventional high-density emitter localization methods that are based on iterative numerical optimization, our WindSTORM uses non-iterative linear deconvolution to decompose overlapping emitters and retrieve their precise locations. Through numerical simulation and biological experiments, we demonstrate that WindSTORM achieves real-time image processing of high-throughput nanoscopy on a GPU device and maintain high accuracy and fidelity even in the presence of high non-uniform background in various biological samples.

## Results

### WindSTORM

The central task of a high-density emitter localization method is to identify and localize overlapping emitters. Our WindSTORM achieves robust and high-speed high-density emitter localization via two major steps: (1) overlapping emitter decomposition by background removal and inverse deconvolution with frequency truncation to identify overlapping emitters; and (2) emitter localization via surrounding emitter deduction and central emitter recovery followed by gradient fitting to precisely localize emitters.

The complexity of overlapping emitter model is the main hurdle for the development of a simple solution to localize overlapping emitters in the high-density emitter scenarios. All existing high-density emitter localization methods rely on numerical optimization with massive iterations to get approximate estimation of emitter positions^7,11^. Non-iterative linear deconvolution methods (e.g., inverse filtering) are potentially attractive approaches to decompose overlapping emitters, but often suffers from serious limitations: background noise often introduces significant artifacts, limiting its fidelity and its ability to accurately decompose the overlapping emitters.

WindSTORM overcomes these limitations and uses a computationally simple non-iterative high-density emitter localization method. As illustrated in Fig. 1, we first perform background subtraction based on extreme value-based emitter recovery^18^ to minimize artifacts in linear deconvolution, followed by non-iterative inverse filtering and frequency truncation to decompose the overlapping emitters (see Supplementary Note). Note that all truncated spatial frequencies are to remove the noise that much higher than the cutoff spatial frequency determined by the diffraction-limited resolution of the optical system, as indicated by Supplementary Fig. S1. Then the overlapping emitters can be easily identified by finding their local maxima. Last, for subsequent emitter localization, we apply the surrounding emitter subtraction^19^ (see Supplementary Fig. S2 and Supplementary Note) to recover the central emitter within each region of interest, which enables us to use a simple algebraic algorithm of gradient fitting^20–22^ for fast emitter localization. As all steps are based on non-iterative operations, we significantly improve the computational speed of WindSTORM on both central processing unit (CPU) and graphics processing unit (GPU), while maintaining a high accuracy for image reconstruction, even in the presence of non-uniform and high background with weak fluorescent emitters.

**Figure 1.**
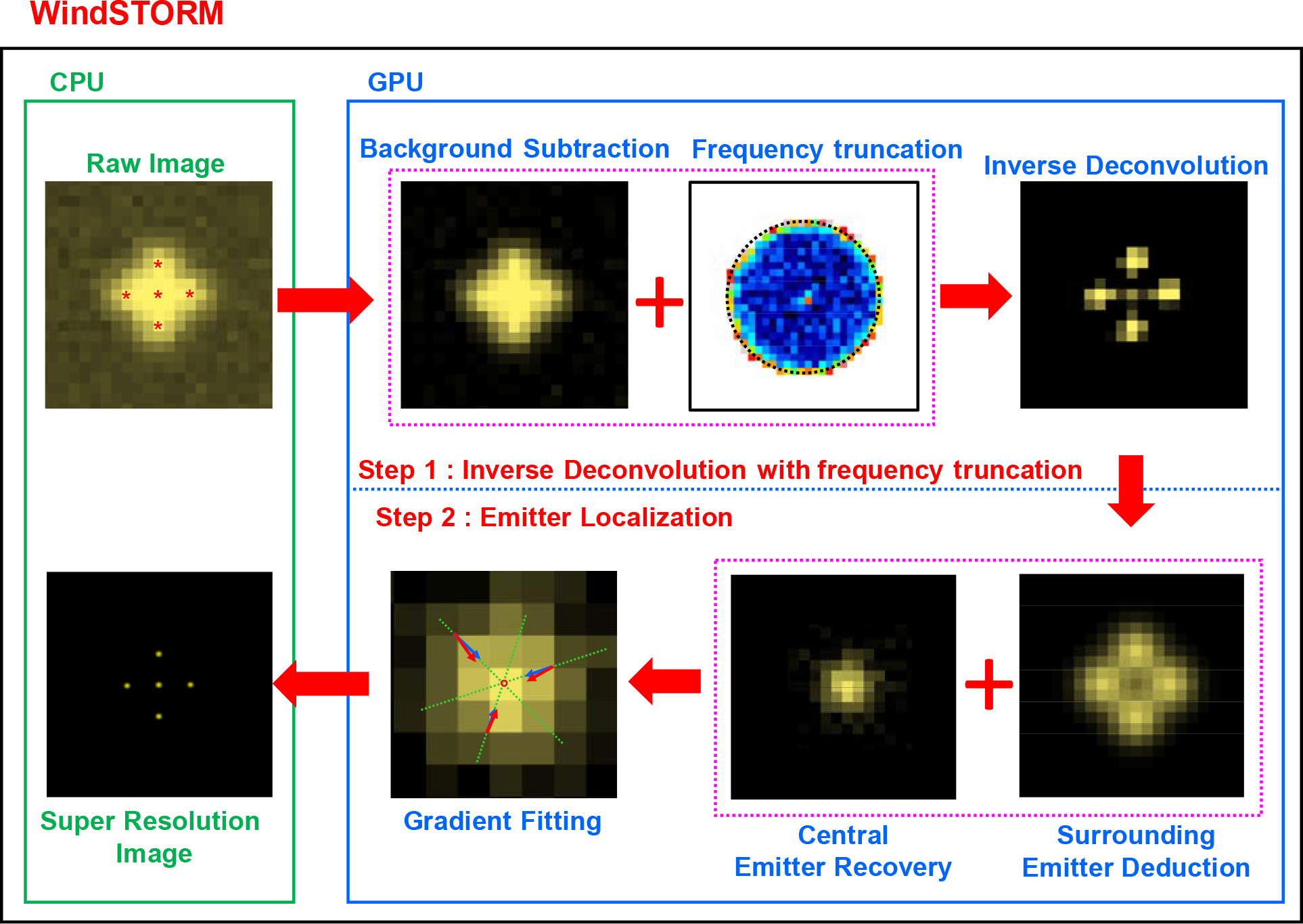
The workflow of high-speed high-density localization method – WindSTORM. In our GPU implementation, all steps (inverse deconvolution and emitter localization) were executed on GPU.

### Numerical simulations

We first benchmarked the performance of WindSTORM against three conventional approaches - a compressed sensing based approach (implemented in FALCON)^23^, a multi-emitter fitting algorithm (implemented in (3D)DAOSTORM^24,25^) and a single-emitter fitting algorithm (implemented in ThunderSTORM^26^) using simulated dataset with a wide range of emitter densities (0.1-6 emitters/μm^2^) in the presence of uniform background. The point spread function (PSF) of each emitter is modeled with the classical Airy pattern derived from the diffraction theory^18^ (see Supplementary Fig. S3a), and the emitter intensity is modeled with a log-normal distribution^23^ (see Supplementary Fig. S3b). The PSF used in WindSTORM is also shown in Supplementary Fig. S3a as a comparison. Figure 2 quantifies the performance of emitter localization using emitter recall rate, root mean square error (RMSE) and false positive rate in three scenarios - high signal-to-noise ratio (SNR) (bright fluorophores such as fluorescent dyes), low SNR and ultra-low SNR (weak fluorophores such as fluorescent proteins). For all three scenarios, WindSTORM shows similar recall rate and localization accuracy to those of FALCON and 3D-DAOSTORM, and significantly outperform the single emitter fitting based ThunderSTORM. Figure 3 shows the comparison of the computational time of these four methods for various emitter densities on both CPU and GPU. At high density scenarios (e.g., 5 emitters/μm^2^), WindSTORM is about one-order-of-magnitude faster than ThunderSTORM and 3D-DAOSTORM and two-order-of-magnitude faster than FALCON on CPU; and is three-order-of-magnitude faster than FALCON when implemented on GPU.

**Figure 2.**
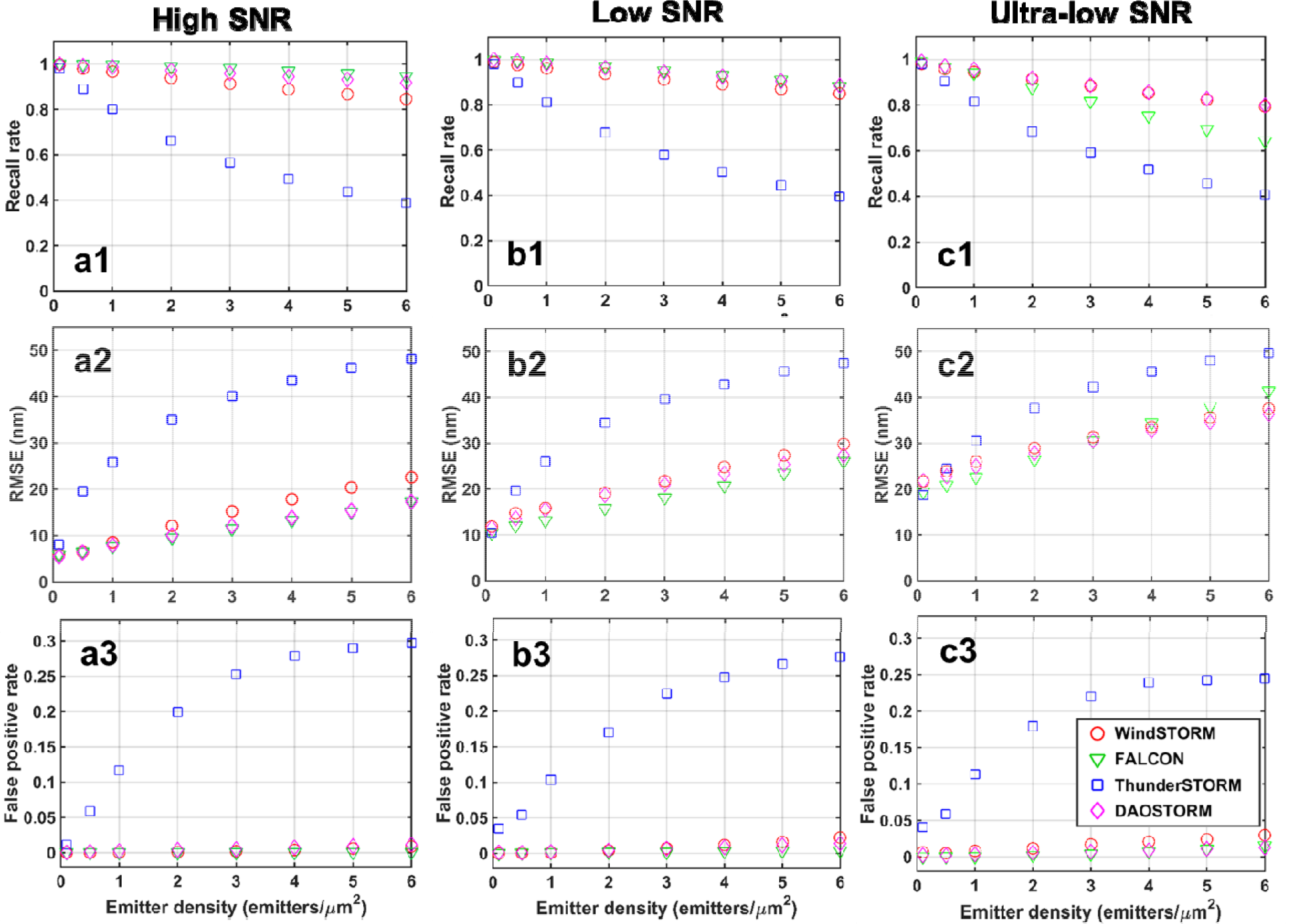
The quantified (**a1-c1**) recall rate, (**a2-c2**) root mean square error (RMSE) and (**a3-c3**) false positive rates for four algorithms – WindSTORM, FALCON, DAOSTORM and ThunderSTORM for various emitter densities using simulated dataset. The heterogeneous emitter intensity is modeled with a lognormal distribution with mean and standard deviation of 5000 and 2000 photons for the high SNR case (**a1-a3**), 1000 and 400 photons for the low SNR case (**b1-b3**) and 350 and 70 photons for the ultra-low SNR case (**c1-c3**).

**Figure 3.**
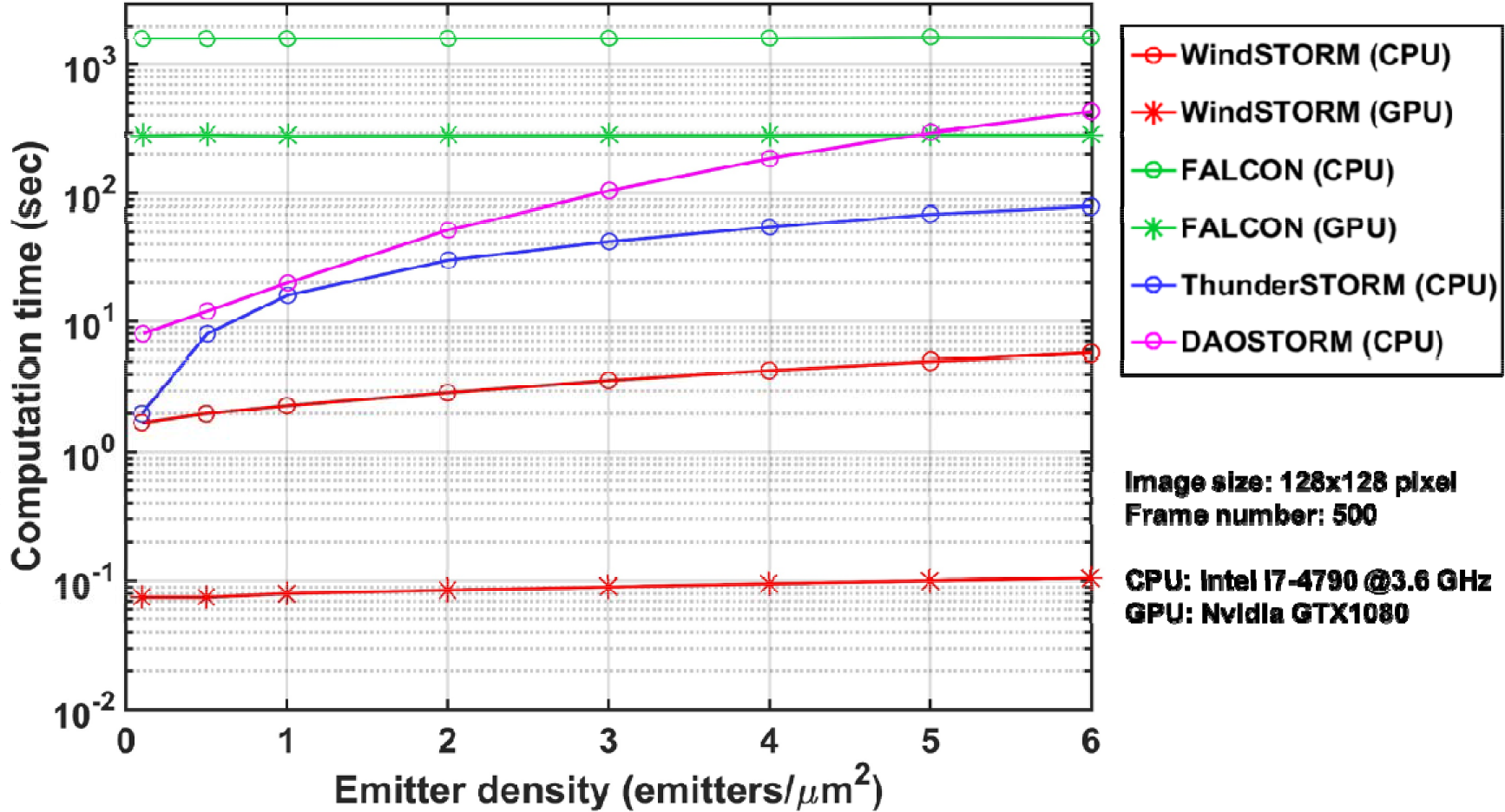
The computation time of image reconstruction by WindSTORM, FALCON, 3D-DAOSTORM and ThunderSTORM on both CPU and GPU for various emitter densities (the same datasets used in Fig. 2).

Next, we evaluated the performance of WindSTORM in a more complex scenario using simulated dataset with dense emitters (5 emitters/μm^2^) of bright emitters (mean μ = 5000; standard deviation σ = 2000 photons) in the presence of non-uniform and high background (ranging from 100 to 1000 photons per pixel per frame) that also undergoes a temporal decay (50%), as shown in Figs. 4a-c. Figures 4d-g compare the ground truth (TRUE) with the reconstructed images by WindSTORM, FALCON and ThunderSTORM, and Fig. 4h compares their localization errors for the recalled emitters. Overall, the single emitter fitting method (ThunderSTORM) suffers from significantly lower recall rate and higher localization error than WindSTORM and FALCON. In the regions with low background (the edge of the image), the images reconstructed by both WindSTORM and FALCON best match with the ground truth. However, in the middle of image where non-uniform background is present (Fig. 4f), there is a significant degradation of localization accuracy of FALCON. Figure 4h shows a larger number of emitters with localization error of >20 nm. Similar results were also seen using simulated dataset of weaker emitters with mean photon counts of 1500 and standard deviation of 500 in the presence of non-uniform background (50-500 photons/pixel/frame), as shown in Supplementary Fig. S4. WindSTORM shows a better accuracy compared to that of FALCON and ThunderSTORM. These results demonstrate the robustness of WindSTORM to accurately reconstruct the super-resolution image in the presence of non-uniform and high background.

**Figure 4.**
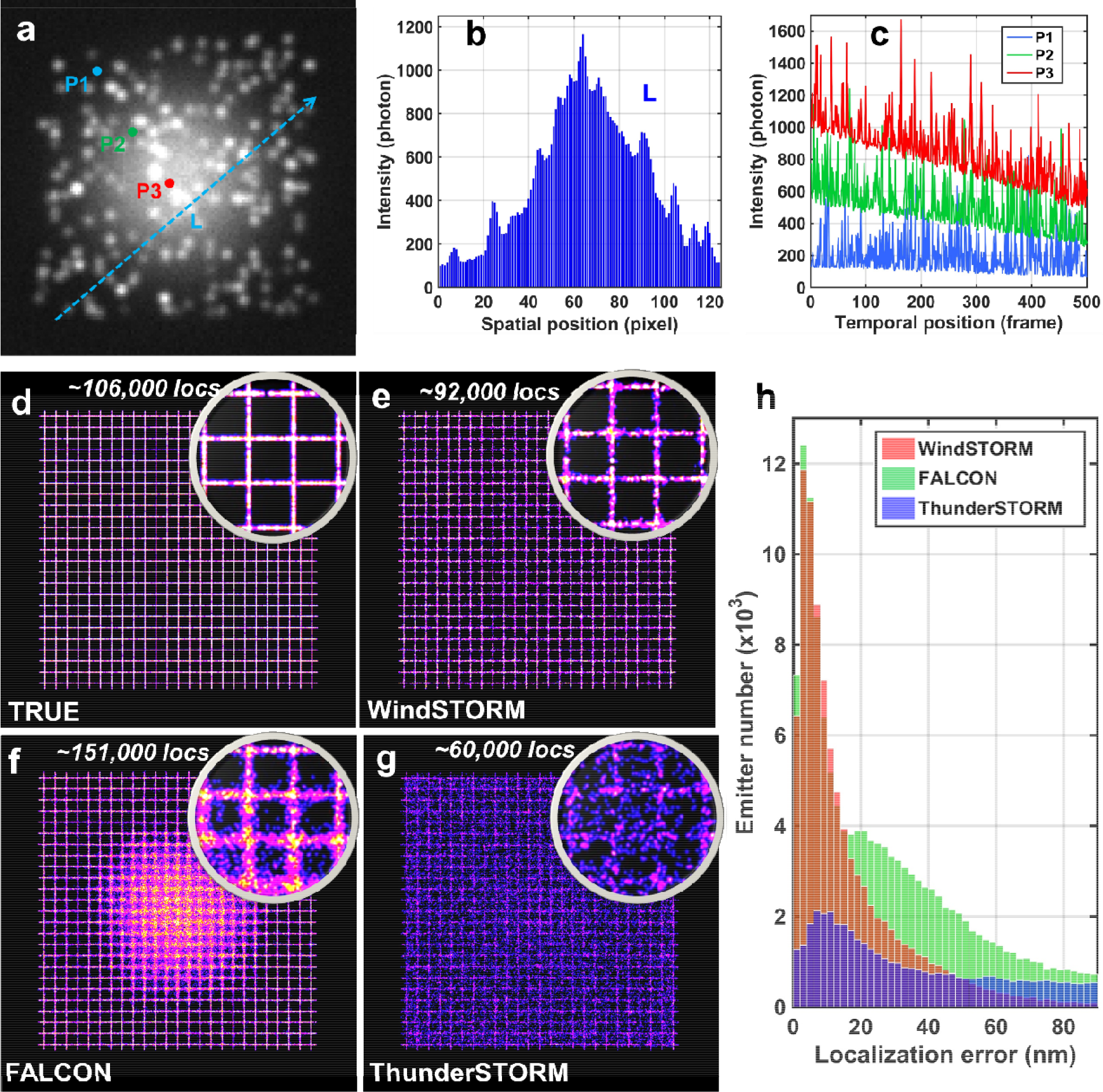
(**a**) A single-frame raw image in the presence of a non-uniform background modeled with Gaussian profile. The background intensity ranges from 100 to 1000 photons. (**b**) The cross-sectional profile of the dashed line (L) shown in (a), which clearly shows a non-uniform background. (**c**) The temporal profile of intensity at 3 locations (P1, P2, P3) in (**a**) that decay linearly over time (∼50% reduction at 500 frames). (**d-g**) The ground truth (**d**) and the super-resolution images reconstructed with WindSTORM (**e**), FALCON (**f**) and ThunderSTORM (**g**). The simulated image size is 128 × 128 pixels, with emitter density of 5 emitters/μm^2^ and mean emitter intensity of 5000 photons for a total of 500 frames and the total number of emitters of ∼1.1×10^5^. The localized emitter by each method is shown on the top of each image. (**h**) The distribution of localization error for the recalled emitters by WindSTORM, FALCON, and ThunderSTORM.

### Experimental dataset of STORM imaging with relatively uniform background

We then assessed WindSTORM using the experimental dataset of dense microtubules labeled with Alexa Fluor 647 from the open-access reference experimental high-density dataset of tubulins^11^ (see Supplementary Fig. S5), as well as the experimental dataset obtained by our custom-built STORM system (Fig. 5). In our experimental dataset, the mean photon count is about 10000 (see Supplementary Fig. S6a). As shown in Figs. 5a-d, despite the highly dense emitters in a single frame (Fig. 5a), the background is relatively low and uniform (Fig. 5b) with small temporal variation (Fig. 5c). The super-resolution images reconstructed by WindSTORM (Fig. 5d) and FALCON (Fig. 5e) show significantly higher image contrast with more recalled emitters (4.6 million by WindSTORM and 6.1 million by FALCON) and higher Fourier ring correlation (FRC) resolution^27^ (48 nm by WindSTORM and 67 nm by FALCON) than those (2.6 million recalled emitters and 80 nm FRC resolution) by ThunderSTORM. The results from ThunderSTORM also confirm that the emitter density used here is indeed a high-density scenario where single-emitter fitting method suffers from low recall rate and localization accuracy. Importantly, WindSTORM reconstructs the entire image in 11 minutes on CPU (single-core), and 8.5 seconds on GPU, which are much faster than the speed of conventional high-density and low-density localization algorithm FALCON on CPU and GPU. As WindSTORM executed on GPU is faster than the image acquisition time of 100 seconds, it realizes online image reconstruction for high-density emitters.

**Figure 5.**
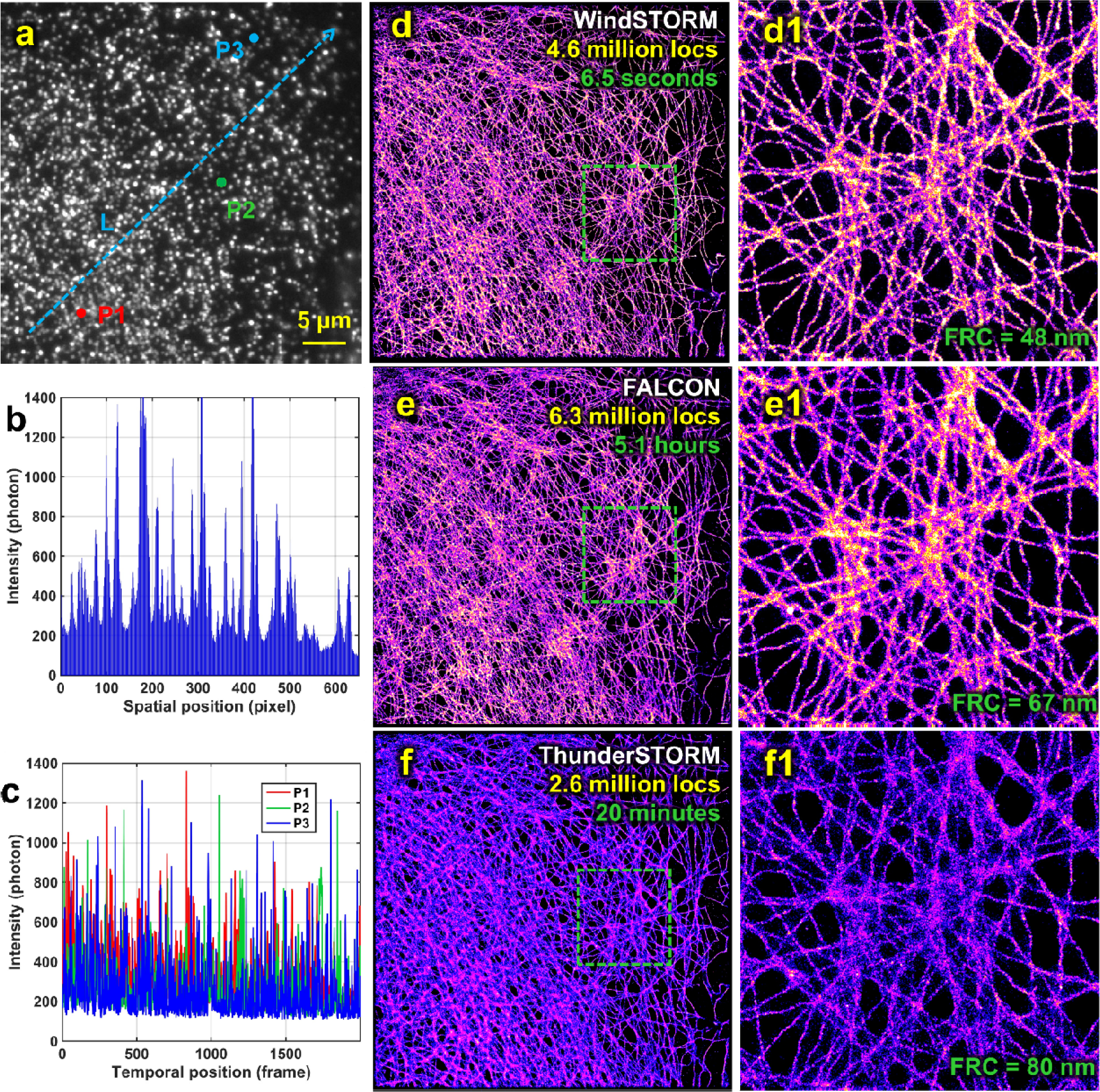
(**a**) A single-frame raw image of tubulin labeled with Alexa 647 in COS-7 cells. The image size is 512 × 512 pixels, with pixel size of 81 nm. (**b**) The cross-sectional profile of the dashed line (L) shown in (a), which clearly shows a non-uniform background. (**c**) The temporal profile of intensity at 3 locations (P1, P2, P3) in (**a**). (**d-f**) Super-resolution images reconstructed by WindSTORM, FALCON and ThunderSTORM, where the number of recalled emitters (locs) and computational time on the same computation platform are shown at the upper right corner. The zoomed regions in the green box are shown in (d1-d1). The FRC resolution is shown at the lower right corner. A total of 2000 frames were used for image reconstruction.

### Experimental dataset of PALM imaging with non-uniform background

We also tested another scenario of PALM imaging of microtubules labeled with photoactivatable fluorescent protein mEoS3.2 which presents non-uniform background. The mean photon count is about 1000 (Supplementary Fig. S6b). As shown in Figs. 6a-c, the intensity profile along the dashed line (L) in (Figs. 6a-b) shows a highly non-uniform distribution, which also undergoes a significant decay over time (Fig. 6c). Figures 6(d-g) compare the diffraction-limited wide-field image (Fig. 6d) with the corresponding super-resolution images reconstructed by WindSTORM (Fig. 6d), FALCON (Fig. 6e) and ThunderSTORM (Fig. 6f). Overall, the images reconstructed by WindSTORM and FALCON exhibit higher image contrast with more recalled emitters compared to that by ThunderSTORM, which confirms the dense emitter scenario. The zoomed images (Figs. 6(d1-g1)) in the green box 1 of Figs. 6(d-g) clearly show the ripple-like artifacts in the image reconstructed by FALCON (Fig. 6f1) that are not present in the diffraction-limited wide-field image (Fig. 6d1), as indicated by the green circles; while the image reconstructed by WindSTORM shows closer resemblance to the diffraction-limited image. Figures 6(d2-g2) shows that the image by FALCON (Fig. 6e2) exhibits the similar ripple-like structure in the circled region as those in Fig. 6e1, likely an artifact, which is not present in the images reconstructed by WindSTORM and ThunderSTORM. Although the ground truth is unknown, our direct comparison between the diffraction-limited and super-resolution images of the same sample has been a well-accepted approach to identify image artifacts in super-resolved images ^28^. Further, WindSTORM takes 79 seconds to reconstruct the image on CPU and 1.8 seconds on GPU, about 2-3 orders of magnitude faster than FALCON on CPU and GPU. This result demonstrates that WindSTORM is not only significantly faster than other high-density localization algorithms, but also more robust to localize weaker emitters (e.g., fluorescent proteins) in the presence of non-uniform background, compared to other methods. Additional experimental data on PALM imaging of other sub-cellular organelles (vimentin and endoplasmic reticulum) are shown in Supplementary Figs. S8–S9.

**Figure 6.**
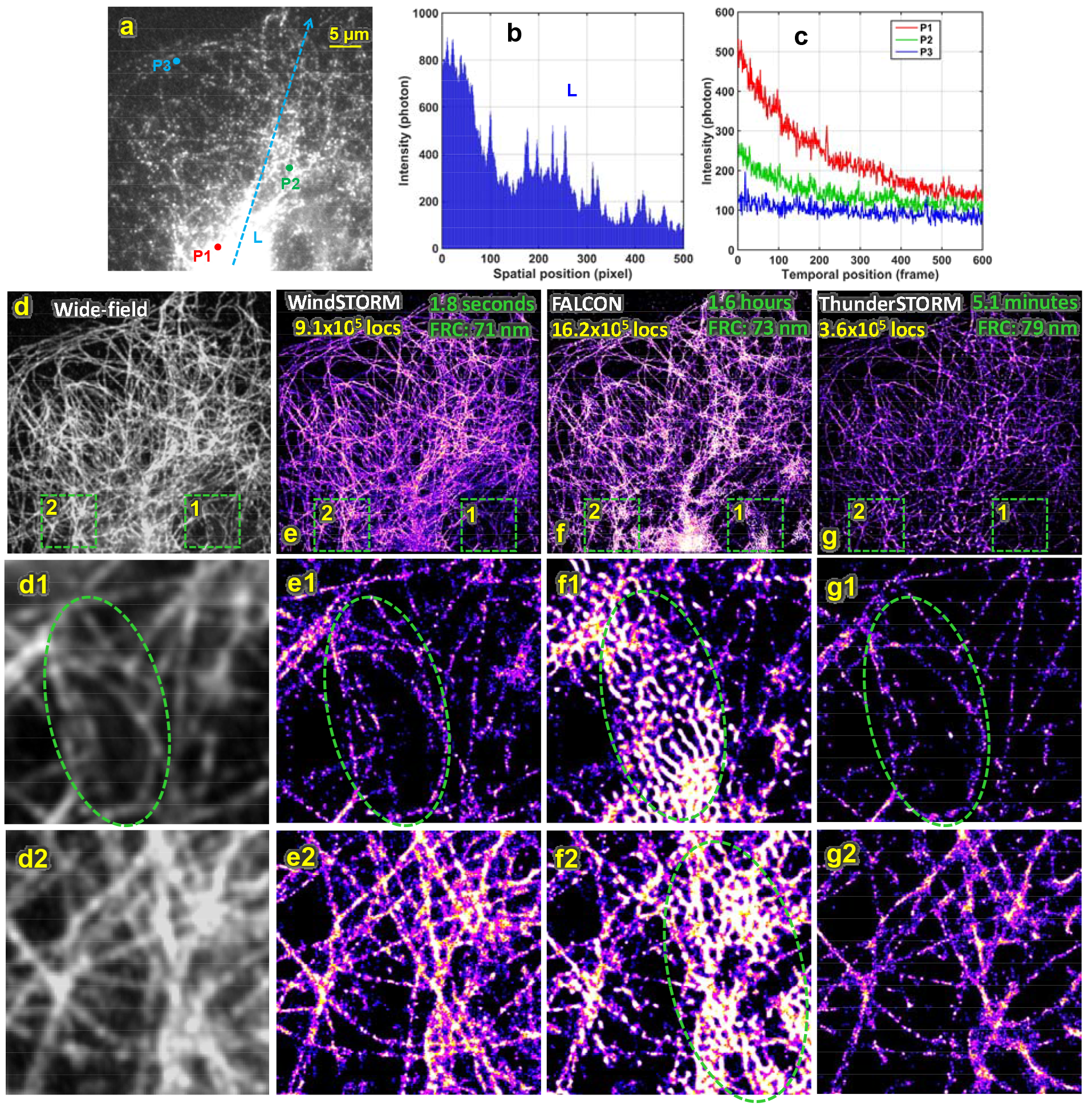
(**a**) A single-frame raw image of tubulin labeled with mEoS3.2 in COS-7 cells which presents non-uniform background. The image size is 512 × 512 pixels and the pixel size on the sample is 81nm. (**b**) The cross-sectional profile of the dashed line (L) shown in (a), which clearly shows a non-uniform background. (**c**) The temporal profile of intensity at 3 locations (P1, P2, P3) in (**a**). (**d-g**) The diffraction-limited wide-field image (**d**) and the corresponding super-resolution images reconstructed by WindSTORM (**e**), FALCON (**f**) and ThunderSTORM (**g**). The number of recalled emitters (locs), the FRC resolution and computational time on the same computing platform are shown at the top of (e-g). The zoomed region in the green boxes 1 and 2 are shown in (d1-g1) and (d2-g2), respectively. A total of 600 frames were used for image reconstruction.

### Experimental dataset of bright emitters with non-uniform and high background

We further evaluated another challenging scenario of imaging densely packed chromatin structure in a tissue section, which often presents heterogeneous strong background due to tissue autofluorescence and stronger scattering. As shown in the representative single-frame raw image (Fig. 7a) of histone mark acetylated H4 (H4Ac) and the intensity profile (Fig. 7c), there is a strong and non-uniform background that also undergoes slow decay over time (Fig. 7d). The super-resolution image reconstructed by WindSTORM shows the distinct clusters formed by H4Ac, which is similar to the previously reported spatially segregated nanoclusters in cultured cells via STORM imaging in the presence of low and uniform background^29^. While the super-resolution images reconstructed by FALCON and ThunderSTORM do not clearly show distinct clusters, but more diffuse structures, likely due to compromised resolution. It is also reflected in the best FRC resolution in the reconstructed image by WindSTORM. In addition, the images reconstructed by FALCON and ThunderSTORM also exhibit background-induced artifacts that are not present in the wide-field image (the region marked by the green circle). To further confirm the presence of clustered structures of H4Ac in the intestinal tissue section, we performed STORM imaging of H4Ac on an ultrathin frozen tissue section, where the background is rather low with sparse emitters. The similar cluster-like structure was also observed in the reconstructed super-resolution image (see Supplementary Fig. S9). Overall, WindSTORM takes 110 seconds to reconstruct the image on CPU and only 6.3 seconds on GPU, also much faster than that of FALCON on CPU and GPU. This result further supports that WindSTORM achieves the speed for online image reconstruction, while maintaining robust performance for high-density emitters in the presence of non-uniform and high background.

**Figure 7.**
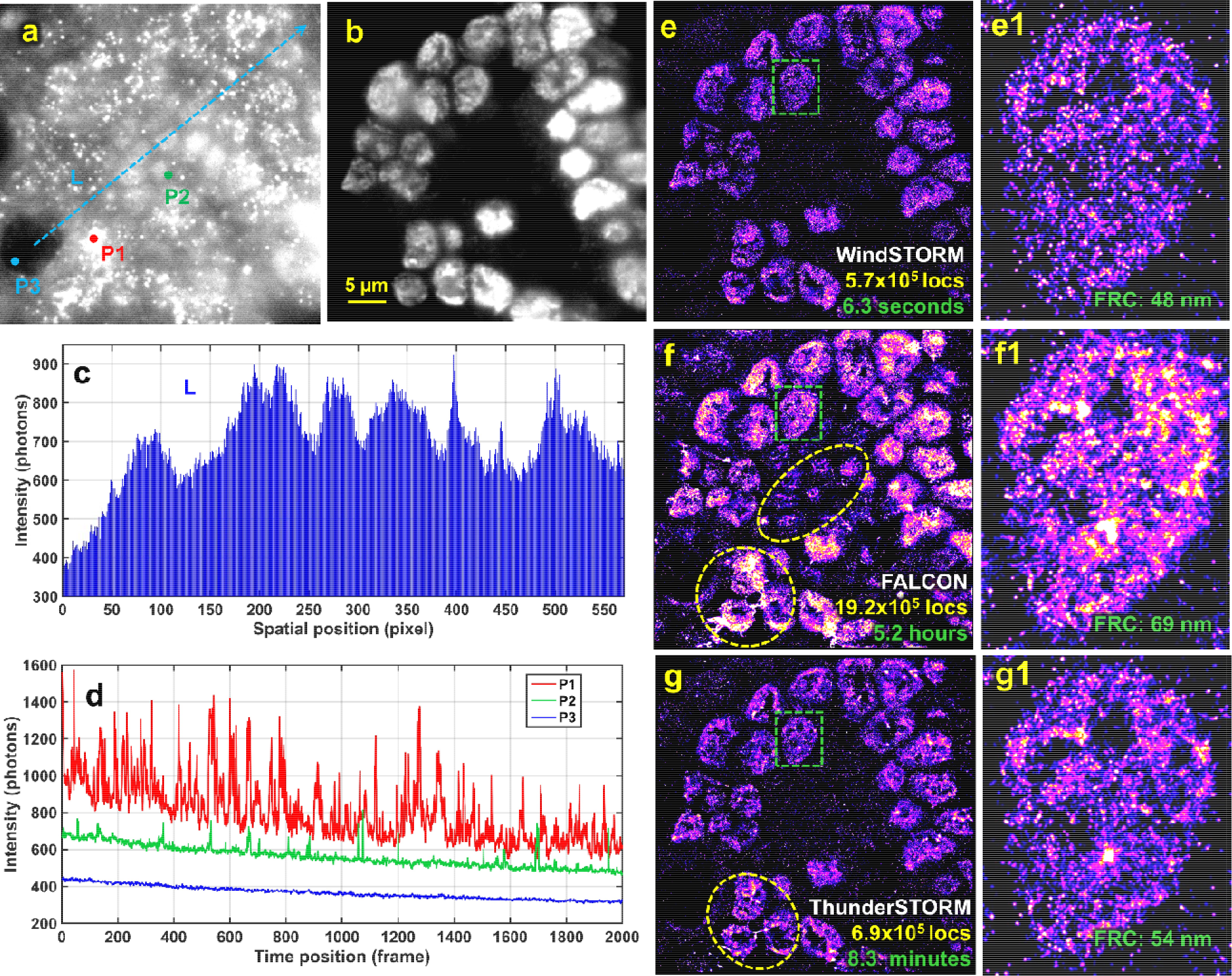
(**a**) A single-frame raw image and (**b**) the corresponding wide-field image of histone mark of acetylated H4 (H4Ac) labeled with Alexa 647 in a mouse intestinal tissue section. The tissue section was cut from formalin-fixed, paraffin-embedded (FFPE) tissue block and the tissue thickness after staining was estimated to be ∼5 μm. The image size is 512 × 512 pixels and the pixel size on the sample is 81nm. (**c**) Cross-sectional intensity profile of the dashed line (L) shown in (a), which shows non-uniform distribution. (**d**) The temporal profile of intensity at 3 locations (P1, P2, P3) in (**a**). (**e-g**) Super-resolution images reconstructed by WindSTORM, FALCON and ThunderSTORM, where number of recalled emitters and computational time on the same computing platform are shown at the bottom right corner. The zoomed region in the green box is shown in (d1-g1) and the FRC resolution is shown at the lower right corner. A total of 2000 frames were used for image reconstruction.

## Discussion

As the next-generation super-resolution microscopy is moving toward high throughput, it can routinely generate a huge dataset (tens of Gigabytes data per minute), where the image reconstruction would take weeks using conventional high-density localization algorithms. We demonstrate a simple non-iterative high-density emitter localization method—WindSTORM—to achieve high-speed and robust image processing for high-density super-resolution localization nanoscopy. Compared to conventional high-density emitter localization methods, WindSTORM significantly improves the data processing speed by ∼2-3 orders of magnitude and remain robust for heterogeneous background; when executed on GPU, the real-time image reconstruction can be realized. Besides the significantly faster speed, WindSTORM also achieves high localization accuracy for a wide range of biological samples and image characteristics, especially for weaker emitters, non-uniform and high background, where conventional high-density localization methods often suffer from poor localization accuracy and significant image artifacts.

There are two precautions that the users should take when using WindSTORM. First, as WindSTORM is based on fast Fourier transform, the sampling rate of the optical imaging system needs to be adjusted to exceed the Nyquist rate (standard deviation (σ) of the PSF σ_PSF_ ≈ 1 pixel). In our experiments, we tested σ_PSF_ of 1.4 pixels (Supplementary Fig. S5), 1.5 pixels (Fig. 6) and 1.9 pixels (Fig. 5), and all of these over-sampling conditions perform well. An under-sampling condition may compromise the performance of WindSTORM. We recommend σ_PSF_ of 1.2∼2.0 pixels to balance sampling rate and SNR. Second, the background estimation algorithm used in WindSTORM is developed based on shot noise model (for sCMOS camera) for temporally slowly-varying background. If EMCCD camera is used where the excess noise caused by electron multiplication cannot be ignored, the background correction model needs to be adjusted to account for excess noise^18^. However, for high-throughput nanoscopy, sCMOS camera is highly preferable due to its large field of view and fast frame rate.

In WindSTORM, the width of the PSF needs to be specified as an input parameter. Although this parameter can be easily measured from the experimental data ^30^, we also explored the scenarios when there is a significant mismatch in PSF width between that used in the inverse deconvolution (WindSTORM) and the actual dataset. Given the Airy-shaped PSF of the optical system with a width of σ=1.5 pixels, we tested the performance of WindSTORM for mismatched PSF not only in shape (Gaussian-shaped PSF used in deconvolution) but also in PSF width. As shown in Figs. S11(a-e), the mismatched PSF width (±0.2 pixel) results in ∼1-2% reduction in emitter recall rate and ∼3-4 nm reduction in localization accuracy. Similar results are also found in the experimental dataset, as shown in Supplementary Fig. S11 (f-i). Overall, for mismatched PSF between WindSTORM and actual dataset, the performance of WindSTORM is slightly affected.

In many biological applications, the mixed scenarios of high-density and sparse emitters can be present within the same sample, so the choice of faster image reconstruction speed with single-emitter algorithms may compromise the image resolution. WindSTORM overcomes this limitation, with superior speed, accuracy, emitter recall rate and robustness for various imaging conditions compared to conventional localization algorithms. It also eliminates the need for the “expert” users to select the proper localization algorithms (single-emitter or high-density) based on the emitter density in the raw image and reduces the barrier for the widespread use of super-resolution localization microscopy.

## Methods

### Implementation of WindSTORM in GPU

Our WindSTORM, including inverse deconvolution, emitter identification and extraction, and emitter localization steps, is fully implemented in GPU to realize online image processing. The inverse deconvolution (Fourier transform) in Step 1 is implemented based on the cuFFT library of CUDA, and emitter localization in Step 2 is implemented according to the literatures^31^, where every emitter is assigned to one thread for parallel processing. Emitter identification and extraction steps, which were often implemented as serial steps in conventional GPU implementation^31^, is also fully parallelized in our implementation to boost the image processing speed. The parallelization of these steps simultaneously eliminates the need for frequent rate-limiting data exchange between CPU and GPU, which can be very time-consuming for large dataset. And thus, our WindSTORM can be robustly executed in GPU and consumes little resources like CPU and memory on personal computers for high-throughput nanoscopy even with huge dataset.

### Numerical Simulation

We simulated a series of image sets with a wide range of emitter densities from 0.1 emitters/μm^2^ to 6 emitters/μm^2^ commonly seen in super-resolution localization microscopy. Given that the localization error inevitably increases with emitter density regardless of localization algorithms, we limit our emitter density to be no higher than 5-6 emitters/μm^2^ to ensure a satisfactory spatial resolution of ∼50-60 nm. Further increase of emitter density will come at the expense of localization accuracy^5^. The image size was set to be 128×128 pixels with a pixel size of 65 nm to mimic a super-resolution microscopy with 100X objective lens (NA=1.49) and sCMOS cameras (pixel size: 6.5 μm), and the emitters were randomly distributed in the image. For each image frame, the fluorescence signal is modeled as a distribution of emitters convolved with an Airy pattern, whose kernel width is set to be 1.5 pixels (∼100 nm). Emitter intensity is set to follow a log-normal distribution, with mean (μ) and standard deviation (σ) of 5000 photons and 2000 photons to mimic bright fluorophores (e.g., Alexa Fluor 647), with μ = 1000 photons and σ = 400 photons to mimic the dimmer fluorescent proteins, and μ = 350 photons and σ = 70 photons to mimic the ultra-low fluorescent intensity (e.g., mEoS3.2). The background per pixel per frame is set to be 50, 20 and 10 photons, respectively, for bright, dim and ultra-low emitter scenarios.

To test the performance of these algorithms under high and non-uniform background, we simulate a dataset with an emitter density of 5 emitters/μm^2^, the emitter intensity follows a log-normal distribution with mean of 5000 photons and standard deviation of 2000 photons, with a Gaussian shaped (sigma = 20 pixels) background at a peak intensity of 1000 photons and an offset of 100 photons added to the image (see Fig. 4). We also performed a similar simulation using mean photon of 1500 and the standard deviation of 500, with a non-uniform background (50∼500 photons with Gaussian shape, see Supplementary Fig. S4).

### Optical setup and image acquisition

The experiments were performed on our home-built super-resolution fluorescence microscopy based upon an Olympus IX71 inverted microscope as described previously^32^. The fluorophore (Alexa Fluor 647) or the fluorescent protein (mEoS3.2) were excited by 642 nm or 560 nm lasers, respectively (VFL-P-1000-642-OEM3 and VFL-P-200-560-OEM1, MPB Communications). The 405 nm laser (DL405-050, CrystaLaser) was used for activation. The laser intensity was controlled by a neutral density filter (NDC-50C-4-A, Thorlabs). The laser beam was expanded by a 10X beam expander (T81-10X, Newport), adjusted for the appropriate field of view. The laser beam was focused onto the rear pupil of a 100X, NA-1.4 oil immersion objective (UPLSAPO 100XO, Olympus) by an achromatic lens. A high oblique angle was used to illuminate the sample. The fluorescent light was collected by the objective, passing through the dichroic mirror (ZT488/640rpc-UF1, Chroma used for STORM imaging and ZET405/488/561/640m for PALM imaging) and band-pass emission filters (ZET488/640m, Chroma for STORM imaging and ET610/75m for PALM imaging), and then focused by the tube lens onto a sCMOS camera (pco.edge 4.2, PCO-Tech) with a pixel size of 81 nm. For STORM imaging, we acquired 2,000 image frames under a 640 nm laser with a power density of 3 kW/cm^2^ and 405 nm laser at a power density of 2.5 W/cm^2^ for activation; for PALM imaging, we acquired 600-1,000 image frames under 561 nm laser at a power density of 2 kW/cm^2^ for excitation and 405 nm laser at a power density of 2.5 W/cm^2^ for activation. The exposure time are all set as 50 milliseconds (ms).

### Sample preparation for STORM imaging

Mouse embryo fibroblast (MEF), COS-7 and U2OS cells were grown at 37°C with 5% CO_2_ and maintained in Dulbecco’s modified Eagle’s medium (DMEM) supplemented with 10% fetal bovine. After growing to about 50% confluency, the cells were extracted or fixed, plated onto a PDL (poly-D-lysine)-coated glass-bottom dish (FD3510, World Precision Instruments) and incubated overnight to let the cells attach to the dish.

For immunofluorescence staining, cells were pre-extracted for 30-60 seconds in 0.5% Triton X-100 (Triton) in BRB80 buffer supplemented with 4 mM EGTA and fixed with Methanol (−20°C) for 10 minutes. Then the fixed samples were washed three times with PBS. The blocking buffer (3% BSA + 0.1% Triton X-100 in PBS) was added to incubate for 1 hour, followed by incubation with primary antibody anti-rabbit α-tubulin (Abcam, ab18251) at 1:300 concentration overnight at 4°C. Then the cells were washed three times with washing buffer. The secondary anti-donkey anti-rabbit antibody (Jackson ImmunoResearch, 121165) conjugated with Alexa Fluor 647 (ThermoFisher, A20106) in blocking buffer at 1:200 concentration was added, then incubated for 2 hours at room temperature and protected from light.

For tissue imaging, a 3 μm-thick intestinal mouse tissue section was obtained from formalin-fixed, paraffin-embedded tissue block and placed on a coverslip. The tissue section was first deparaffinized in xylene, rehydrated and followed by heat-induced antigen retrieval performed in the pre-heated Tris-EDTA buffer solution in microwave oven. The section was then stained with the primary antibody (rabbit polyclonal H4Ac, Cat. 06-598, Millipore) and goat-anti-rabbit secondary antibody conjugated with Alexa Fluor 647 following the same steps as described above.

Immediately before imaging, PBS buffer was replaced with imaging buffer composed of 10% w/v glucose (Sigma-Aldrich), 0.56 mg/mL glucose oxidase (Sigma-Aldrich), 0.17 mg/mL catalase (Sigma-Aldrich), 0.14M β-mercaptoethanol (Sigma-Aldrich) and 2 mM cyclooctatetraene (COT).

### Sample preparation for PALM imaging

The mEos3.2-Tubulin-C-18, mEos3.2-Vimentin-C-18 and mEos3.2-ER-5 were gifts from Michael Davidson (Addgene plasmid # 57484, #57486 and #57455, respectively). Cells were transfected using 2 mL transfection media, 100 μL Opti-MEM (Gibco), 3 μL Lipofectamine2000 (Invitrogen), and 1 μg plasmid per 3.5 cm MatTek glass-bottom dish. After 24 hours of incubation, cells were fixed with 4% PFA in PBS for PALM imaging.

### Image reconstruction

We reconstructed the super-resolution image with a single-emitter fitting method (ThunderSTORM), a multi-emitter fitting method (3D-DAOSTORM) and a compressed sensing-based approach (FALCON), all compared with WindSTORM. The single-emitter maximum likelihood Gaussian function fitting algorithm and wavelet filtering are used in ThunderSTORM, multi-emitter maximum likelihood Gaussian function fitting algorithm is used in 3D-DAOSTORM. The kernel width of the PSF used in WindSTORM is set to 1.5 pixels for simulated dataset and our experimental PALM imaging dataset, 1.9 pixels for our experimental STORM imaging dataset and 1.4 pixels for the open-access experimental dataset (http://bigwww.epfl.ch/smlm/datasets/index.html?p=real-hd). For iterative approaches (FALCON and ThunderSTORM), the above kernel widths of the PSF were used as the initial kernel. The drift correction was performed using cross correlation method provided in ThunderSTORM. In this study, the GPU versions of WindSTORM and FALCON are executed on GPU (GTX1080, Nvidia), the CPU version of WindSTORM, 3D-DAOSTORM, FALCON and ThunderSTORM are executed on a Quad-core CPU (Core i7-4790, Intel).

## Author contributions

H.M. developed WindSTORM, performed the simulation, data analysis and experiments, and wrote the manuscript. J.X. prepared the sample. Y.L. supervised the project and wrote the manuscript.

## Acknowledgements

This work is supported by National Institute of Health R33CA225494 and R01CA185363.

## Data and code availability

The software and datasets for testing are available on the website (https://pitt.box.com/v/WindSTORM) for public use under CC BY-NC license. The password for the software and datasets generated during the current study are available from the corresponding authors on reasonable request.

## Supplementary Note

### WindSTORM

WindSTORM is a high-speed high-density localization method for high-throughput nanoscopy. It is based on two mathematically simple non-iterative steps: (1) overlapping emitter decomposition via inverse deconvolution with windowed (truncated) frequency; and (2) emitter localization for precise localization via surrounding emitter deduction and gradient fitting.

#### 1. Inverse deconvolution

Linear deconvolution has the potential to achieve fast processing of raw images with high-density emitters due to its non-iterative nature and simplicity. But the presence of non-uniform background often introduces significant artifacts, limiting its use to identify and localize the true overlapping emitters. We made two important modifications to improve its performance in super-resolution localization microscopy.

##### 1.1 Background subtraction

The first step is to subtract the background—based on the conceptual framework of our recently developed extreme value-based emitter recovery method (EVER)^2^—from the raw image before linear deconvolution, which is described below.

###### Theoretical basis of EVER

In super-resolution localization microscopy, the acquired signal can be modeled as Poisson distribution (as shown in the blue curve of Supplementary Fig. S10a), as the read noise, dark noise (generally < 0.1 electrons per frame) and the corrected fixed pattern noise can be neglected for the commonly used sCMOS cameras^3–5^ in high-throughput nanoscopy. The composite signal recorded on the camera consists of the fast-changing emitters and the slowly varying background. In an extreme case with ultra-sparse emitter signal, the composite signal is nearly equivalent to the background signal, which can be well estimated by temporal median value^6^. But in the case of dense emitters when the probability of emitter occurrence within an imaging sequence can be more than 50%, the temporal median value is significantly skewed towards the composite signals, resulting in serious over-estimation of the background signal. However, the extreme value of temporal minimum remains relatively stable regardless of the probability of emitter occurrence, suggesting its inherent robustness. To derive the statistics of temporal minimum, we first consider N random variables to model the photon number *K*_1_, *K*_2_,…, *K*_*N*_, that is independently and identically Poisson distributed with the cumulative distribution function given by

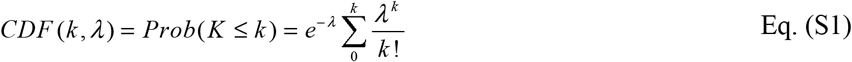

where *λ* is the expected average photon number for each pixel. Then we define a new random variable *K*_min_ = min(*K*_1_, *K*_2_,…, *K*_*N*_) that takes the minimum value among the N Poisson random variables. The cumulative distribution function of this minimum value is

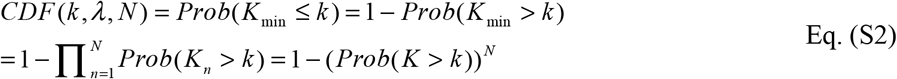

Then the probability mass function (the probability of being the minimum value) is now given by 
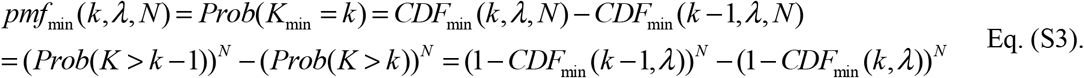

The Eq. S3 assumes uniform background (*λ*) over N (frames). However, for many experimental data, the background also undergoes a slow decay during N frames. To account for such variation, we introduce a decay ratio *R*, and Eq. S3 can be modified to Eq. (4):

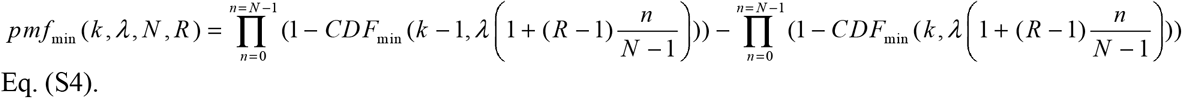

When R = 1, Eq. S4 is reduced to Eq. S3. An accurate estimate of the minimal value requires that the distribution represented by *pmf*_min_ to have a narrow dispersion around its mean (e.g., with a small standard deviation), which can be achieved by applying a spatial mean filter. The distribution can be described by the following Eq. (S5):

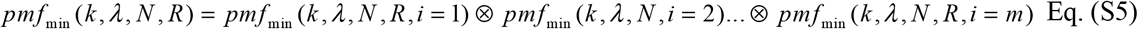

where *m* is the number of pixels being averaged.

Therefore, Equation S5 gives the mathematical solution to describe the probability distribution of temporal minima value, given the expected background value. An example of this distribution is shown in the red curve of Supplementary Fig. S10a (for N = 100 frames, the expected background *λ* = 500 photons, R = 1 and average filter of 3×3), where the dashed red line indicates the expected value (*E*_min_) of the probability distribution of temporal minima.

###### High-speed implementation to estimate background

Based on the above-derived mathematical relationship, one can easily calculate the expected temporal minimum value (*E*_min_) from an image stack, given the expected background (*λ*). But in super-resolution localization microscopy, the inverse problem needs to be solved - estimating the background (*λ*) given the observed temporal minimum value (*E*_min_). The above mathematical relationship (Eq. S5) allows us to create a lookup table for one-to-one mapping between the observed temporal minimum value (*E*_min_) and the expected background (*λ*), as illustrated in Supplementary Fig. S10b.

This lookup table can be large for a wide range of background and index searching of the large look-up table by every thread is highly inefficient in GPU. From the aspect of high-speed computation, to facilitate GPU acceleration, we implemented a simple and high-speed approach by fitting the theoretically-derived look-up table at a given background range (10∼1000 photons) and decay ratio range (1.0∼1.5) that are commonly seen in super-resolution localization microscopy, with a cubic polynomial function 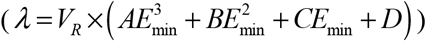. The fitting error for the estimated background is within ± 1.5 photons for the range. Note that, the cubic polynomial fitting function is not the only choice, and other functions such as spline may also be used. The decay factor *V*_*R*_ is calculated for each frame (*f*_*th*_ frame), defined as the average intensity of the non-blinking area (*V*_*f*_) divided by the expected temporal minima (*E*_min_) over the 100-frame sub-stack. To determine the non-blinking area *V*_*f*_, given the slowly-varying non-blinking area and fast-changing emitters, the standard deviation of non-blinking area should be smaller than that of emitters. For Poisson-distributed signals, the mean is equal to the variance. Thus, we defined those pixels with standard deviation (over the 100-frame sub-stack) smaller than two times the square root of *λ*(*R* = 1) (approximated as the mean uniform background) as approximation for the non-blinking area. This estimated background has shown robust performance for a wide range of image characteristics, such as various emitter density, emitter intensity, emitter size, and structured background^2^.

In summary, to perform background estimation, we first segmented the entire raw image stack into a set of sub-stacks along the temporal axis (e.g., 100-frame sub-stack used here), calculated the pixelwise minimum value for each image sub-stack, applied 3×3 average filtering on temporal minima map, and then estimated the background based on the one-to-one mapping between the above-derived expected temporal minimum value and expected background (Supplementary Fig. S10b). Our WindSTORM software already builds in background estimation from the raw image dataset without the need for users’ adjustment. However, if the users use EMCCD camera where the excess noise caused by electron multiplication cannot be ignored, the background correction model needs to be adjusted to account for excess noise.

##### 1.2 Inverse deconvolution and frequency truncation

Next, we perform a linear deconvolution (*F*) based on inverse filtering on the background-corrected image (*I*_*c*_) to recover the overlapping emitters (deconvolved image *D*): *D* = *I*_*c*_ ⊗ *F*. The inverse deconvolution process *F* can be described in the frequency domain (*u*, *v*) as:

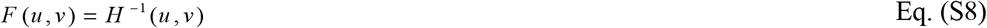
where *H* is the modulation transfer function of point spread function that assumes Gaussian shape.

As shown in Supplementary Fig. S1, after inverse filtering (Figs. S1e1-e2), the high spatial frequency noise dominates. Therefore, we perform truncation on the spatial frequency to remove those high-frequency noise in the deconvolved image in the spatial frequency domain, described by:

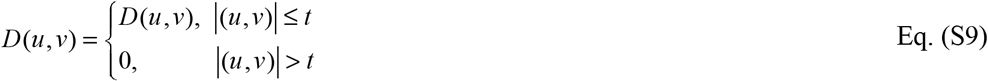

where the frequency cutoff is determined as the point where *D* undergoes a dramatic increase which often occurs at |*H*(*t*)|=0.015×|H(0,0)| for a large intensity range of 150-15000 photons, as shown in Supplementary Fig. S1. Of note, the cutoff spatial frequency determined by the diffraction-limited resolution is still below the truncated frequency, so that such frequency truncation does not reduce the image resolution determined by the optical system.

After the inverse deconvolution, the overlapping emitters are well separated (Figs. S1 (g1-g2)), which can then be identified by finding local maxima, and their approximate positions can be estimated by calculating the center of mass. To identify individual emitters, we used a threshold to avoid artifacts caused by the dim emitters. The threshold is determined as a quarter of the average photon counts of the emitters. This thresholding process is also built into our WindSTORM software, but the user just needs to set the average emitter intensity of each dataset. We suggest that for those who cannot determine this parameter, they can analyze the first frame with single-emitter fitting algorithms (e.g. ThunderSTORM) to get a rough estimation of the emitter intensity.

#### 2. Emitter localization

Next, we perform precise localization of the identified emitters, using our previously developed gradient fitting^7^ to localize the overlapping emitters already identified in the previous step. We previously showed that gradient fitting integrates high speed with high precision. It is an analytical method that runs over two-order-of-magnitude faster than conventional iterative fitting-based algorithm. Its precision outperforms the least square based fitting method and approaches the precision of maximum likelihood estimation (MLE)-based fitting. However, gradient fitting is originally developed for single-emitter localization. To apply it in WindSTORM, we preserved only one central emitter in each region of interest (ROI) by subtracting its surrounding emitters^8^ prior to gradient fitting, as shown in Supplementary Fig. S2. Based on the estimated emitter positions and their relative intensity in the previous step, we can estimate the intensity of each emitter, and subtract its surrounding emitters from the background-corrected raw image (*I*_*c*_), preserving only the central emitter within the ROI for subsequent Gradient fitting.

**Figure S1.**
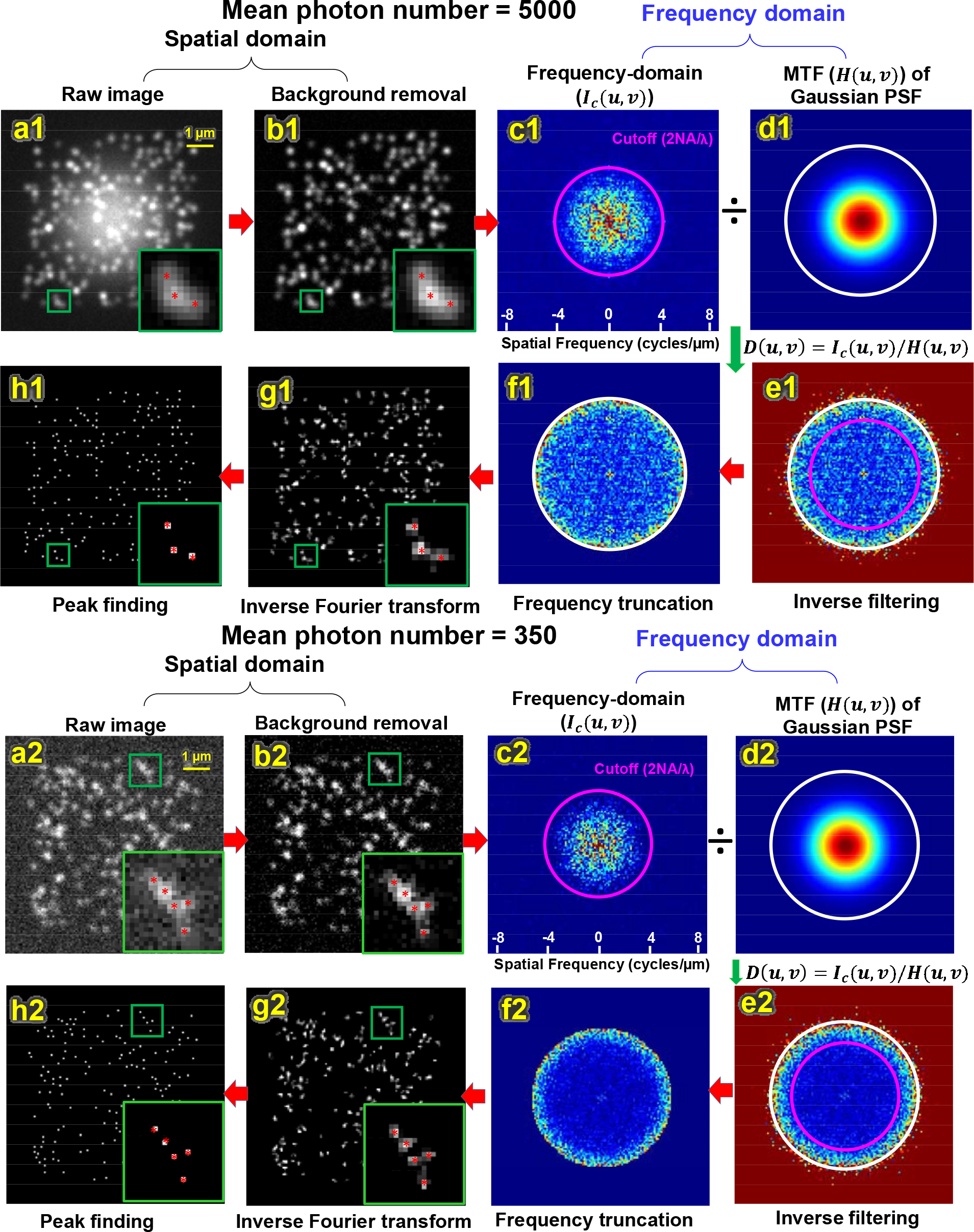
The workflow of inverse deconvolution and frequency truncation for lognormal intensity distribution with (**a1-h1**) 5000 and 2000 (mean and standard deviation) and (**a2-h2**) 350 and 70. (**a1, a2**) Raw single-frame image and (**b1, b2**) the background-corrected image. The insets show the zoomed region of three overlapping emitters and the red stars indicates the central positions of these emitters. (**c1, c2**) The Fourier-transformed image of (b1-b2) (*I*_*c*_(*u*, *v*)) in the spatial frequency domain and the magenta circle indicates the cutoff spatial frequency limited by the resolution of the optical system; (**d1, d2**) the modulation transfer function of a Gaussian PSF (*H* (*u*, *v*)); (**e1, e2**) the deconvolved image after inverse filtering (*D*(*u*, *v*) = *I*_*c*_ (*u*, *v*) / *H* (*u*, *v*)) and the white circle indicates the cutoff spatial frequency for truncation at the threshold. The threshold is defined as the spatial frequency where there is a sharpest change in D. For high and low SNR cases, the cutoff spatial frequency for truncation is approximately the same (*D*_*t*_(*u*_*t*_,*v*_*t*_)  0.015 × *H* (0, 0)); (**f1, f2**) the frequency-domain image after frequency truncation; (**g1, g2**) the deconvolved image in the image plane (after inverse Fourier transform of (e)); the insets show that the overlapping emitters that were hardly identifiable in (a, b) are now well separated. (**h1, h2**) The identified emitters which are used for subsequent emitter localization.

**Figure S2.**
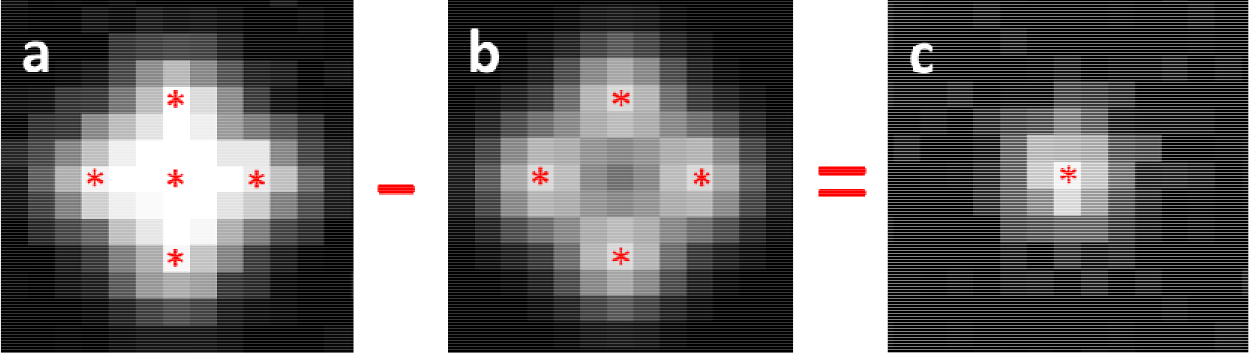
Surrounding emitter deduction to recover the central emitter within the ROI. **(a)** The image at region of interest (ROI); (**b**) surrounding emitters; (**c**) central emitter.

**Figure S3.**
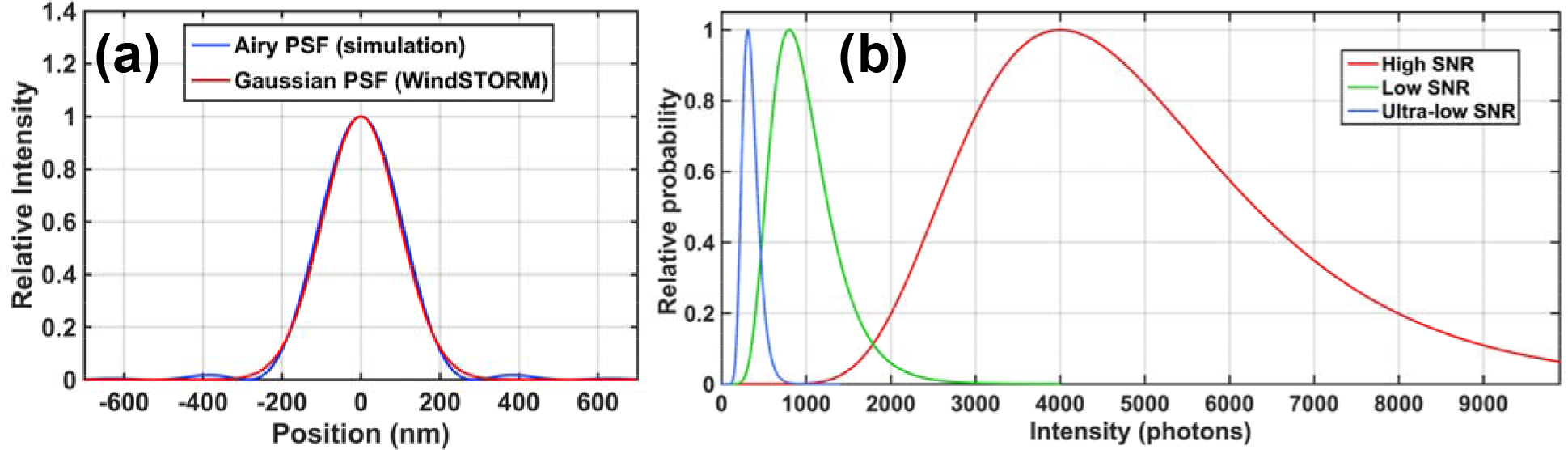
(**a**) The point spread function (PSF) defined by Airy pattern is used for simulation and the Gaussian PSF used for deconvolution in WindSTORM. (**b**) The modeled intensity distribution of emitters that follows a lognormal distribution in the cases of high, low and ultralow SNR.

**Figure S4.**
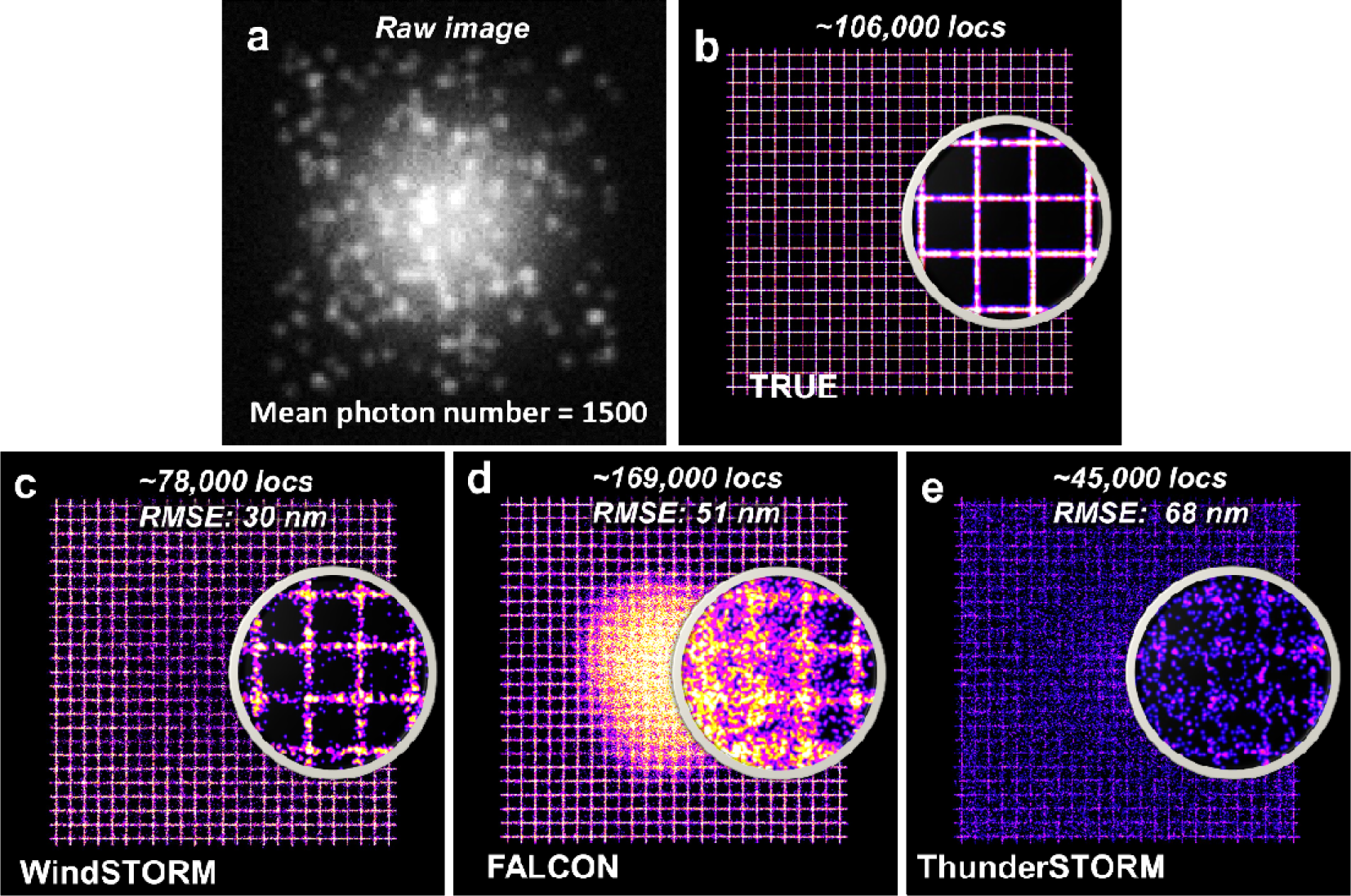
(**a**) A single-frame raw image of simulated dataset with the mean photon number of 1500 (standard deviation is 500 photons) in the presence of a non-uniform background modeled with Gaussian profile. The background intensity ranges from 50 to 500 photons/pixel/frame. (**b-e**) The ground truth (**b**) and the super-resolution images reconstructed with WindSTORM (**c**), FALCON (**d**) and ThunderSTORM (**e**). The simulated image size is 128 × 128 pixels, with emitter density of 5 emitters/μm^2^ and mean emitter intensity of 1500 photons for a total of 500 frames and the total number of emitters of ~1.1×10^5^. The number of localized emitters and root mean square error (RMSE) by each method are shown on the top of each image. The results are consistent with that of Fig. 4 (in the main text). WindSTORM exhibits the best localization accuracy with RMSE of 30 nm, about 21 nm better than FALCON and 38 nm better than ThunderSTORM.

**Figure S5.**
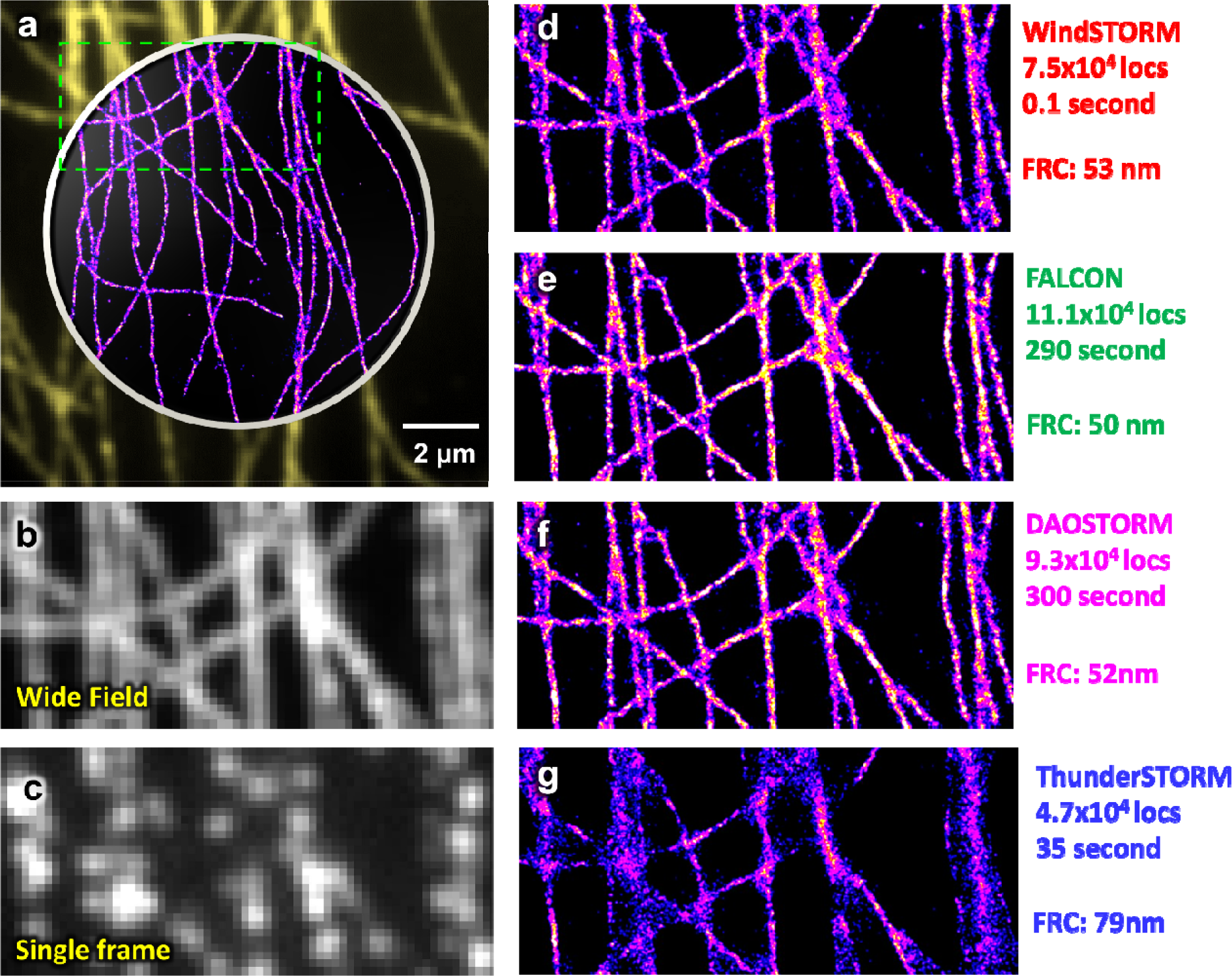
**(a)** The wide-field image and super-resolution image of the open-access experimental high-density localization dataset (available at http://bigwww.epfl.ch/smlm/datasets/index.html?p=real-hd), reconstructed by WindSTORM. **(b-c)** A selected region (green box in a) of wide field and single raw image of fluorescently labeled microtubules. **(d-g)** The final reconstructed super-resolution images from the experimental high-density dataset processed by WindSTORM (GPU), FALCON (GPU), 3D-DAOSTORM (CPU) and ThunderSTORM (CPU). The number of recalled emitters (locs), computational time and FRC resolution of each method are shown on the right-hand side of the image. All high-density localization algorithms (WindSTORM, FALCON, 3D-DAOSTORM) showed better recall rate and FRC resolution than the single-emitter algorithm (ThunderSTORM). WindSTORM shows the fastest times among all methods.

**Figure S6.**
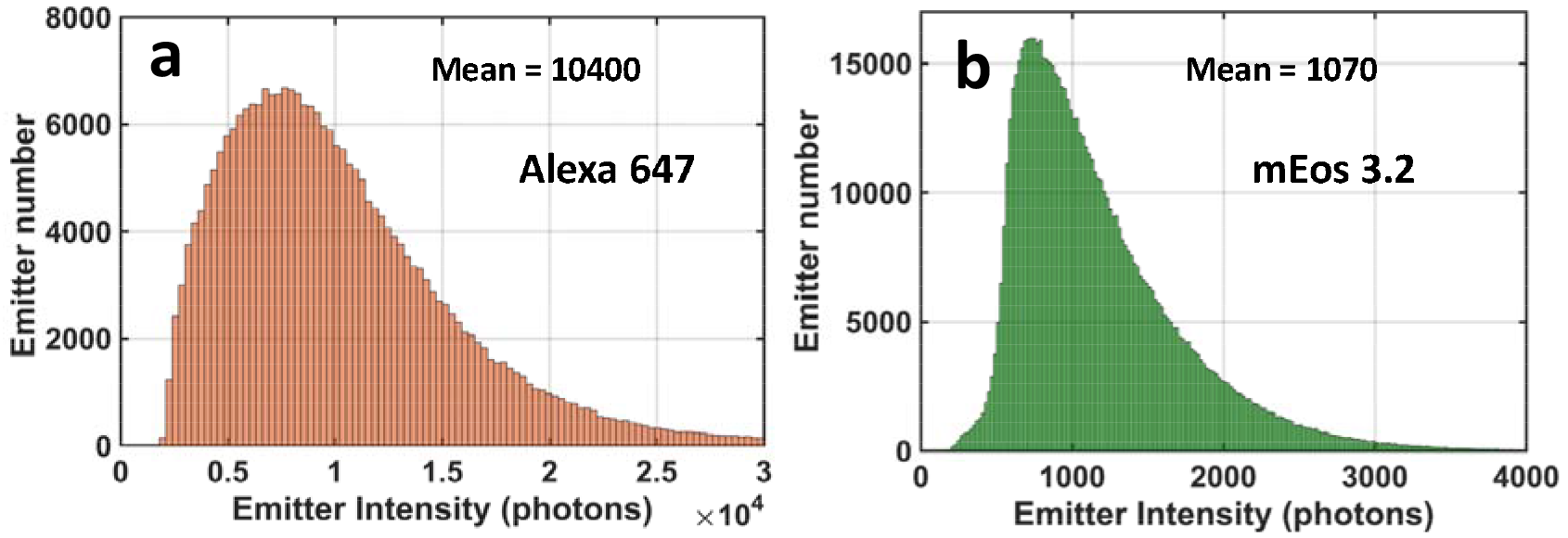
(a) The histogram distribution of emitter intensity (photons) for Alexa 647 and mEoS3.2 used in STORM and PALM imaging of microtubules (Figs. 5 and 6 in the main text).

**Figure S7.**
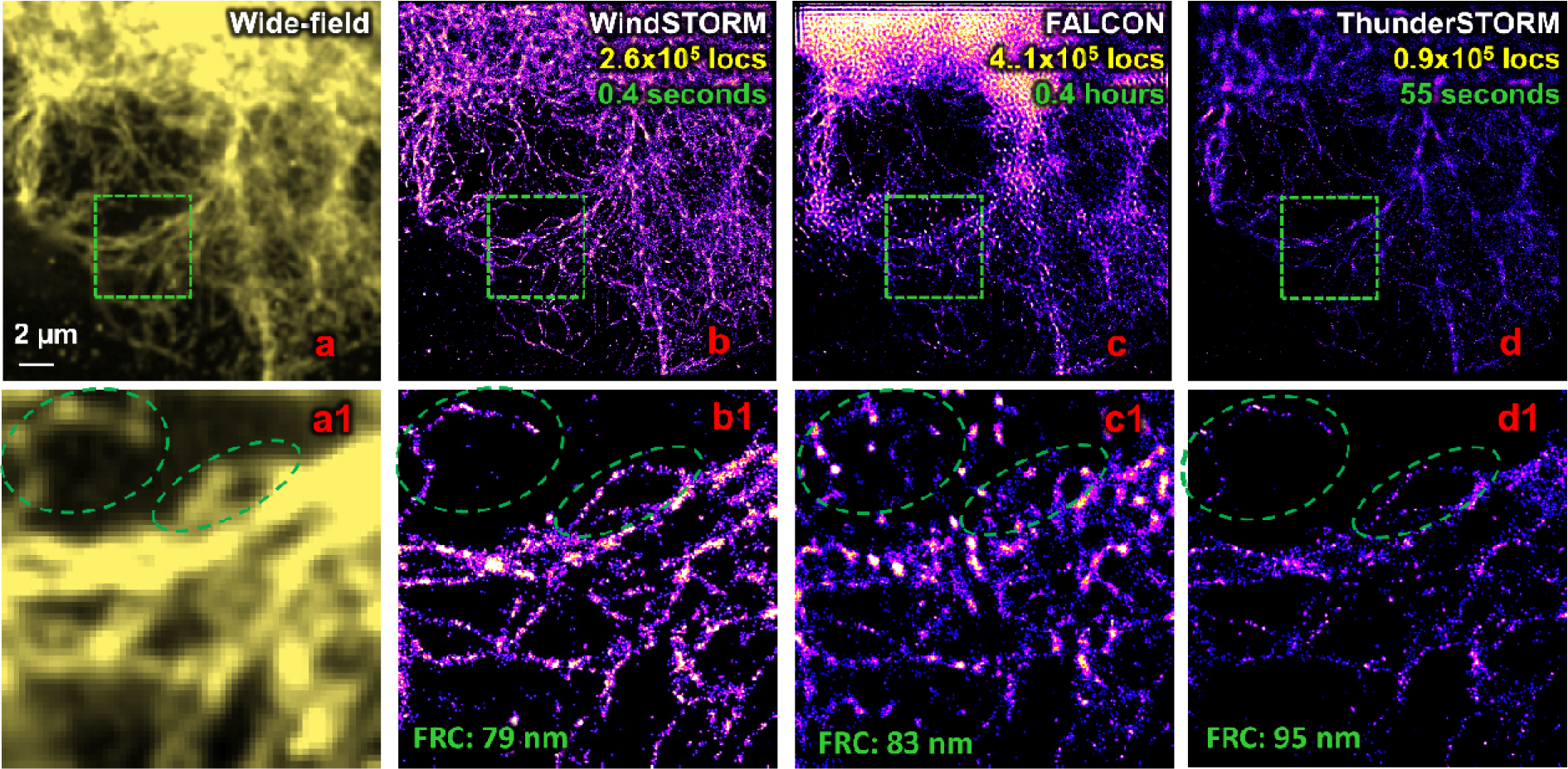
(**a**) The diffraction-limited wide-field image of vimentin (fluorescently labeled with mEoS3.2) in COS-7 cells and corresponding super-resolution images reconstructed by (**b**) WindSTORM (GPU), (**c**) FALCON (GPU) and (**d**) ThunderSTORM (CPU). (**a1-d1**) The zoomed regions of the green boxes in (a-d). The green circles indicate that the structural features from the reconstructed super-resolution image by WindSTORM closely resemble those in the diffraction-limited image (a1), but the reconstructed image by FALCON (c1) shows distinct structural feature, suggesting an image artifact. While the reconstructed image by ThunderSTORM (d1) also shows similar pattern as those of diffraction-limited image, but with less recalled emitters (lower image contrast). The upper right corners of (b-d) show the number of localizations and computation time for WindSTORM (GPU), FALCON (GPU) and ThunderSTORM (CPU).

**Figure S8.**
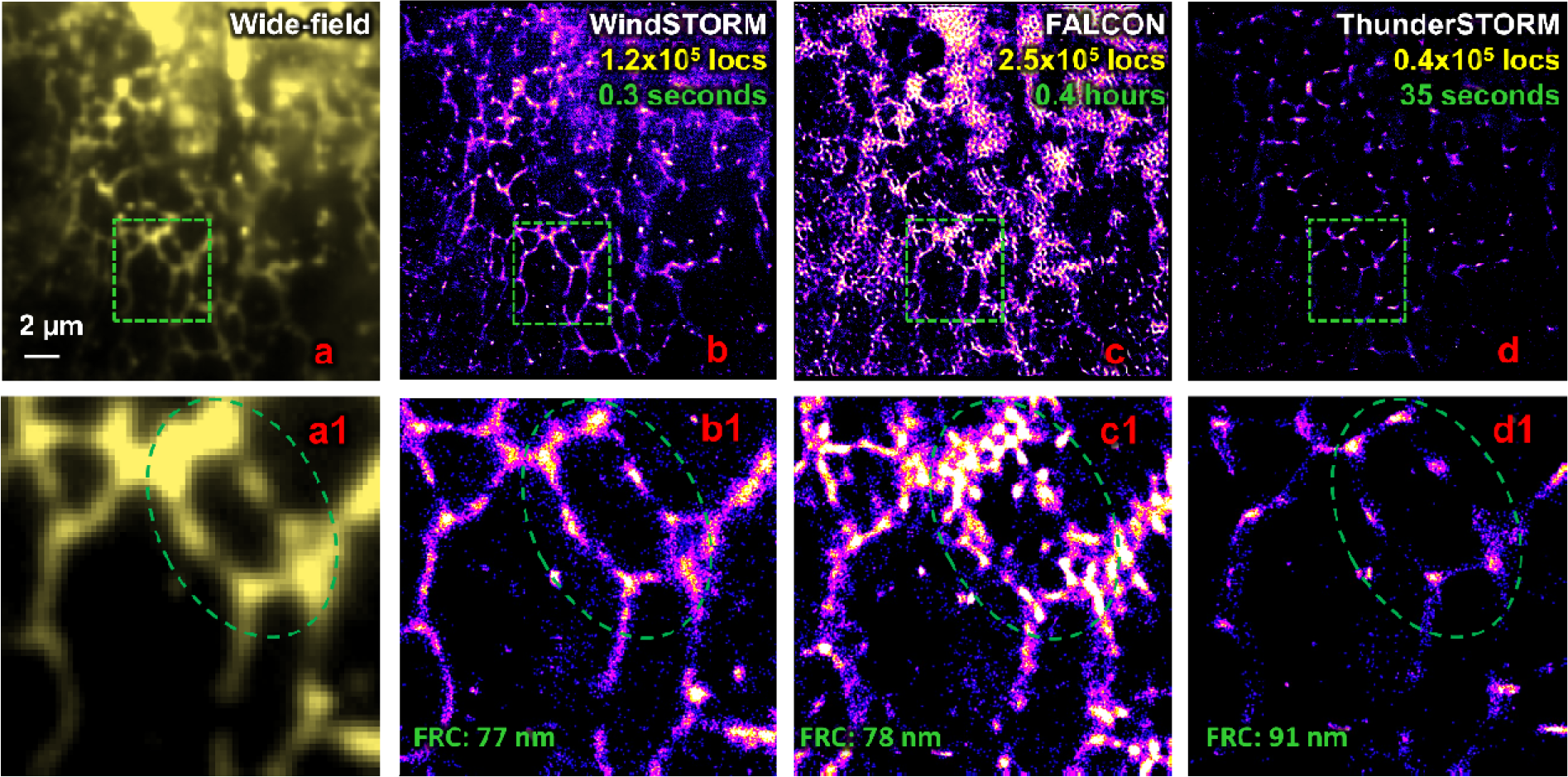
(**a**) The diffraction-limited wide-field image of endoplasmic reticulum (fluorescently labeled with mEoS3.2) in COS-7 cells and corresponding super-resolution images reconstructed by (**b**) WindSTORM, (**c**) FALCON and (**d**) ThunderSTORM. (**a1-d1**) The zoomed regions of the green boxes in (a-d). The green circle indicates that the structural features from the reconstructed super-resolution image by WindSTORM closely resembles the diffraction-limited image (a1), while the reconstructed super-resolution image by FALCON (c1) shows distinct structural feature, suggesting an image artifact. While the reconstructed image by ThunderSTORM (d1) also shows similar pattern as that of diffraction-limited image, but with less continuity and less recalled emitters (lower image contrast). The upper right corners of (b-d) show the number of localizations and computation time for WindSTORM (GPU), FALCON (GPU) and ThunderSTORM (CPU).

**Figure S9.**
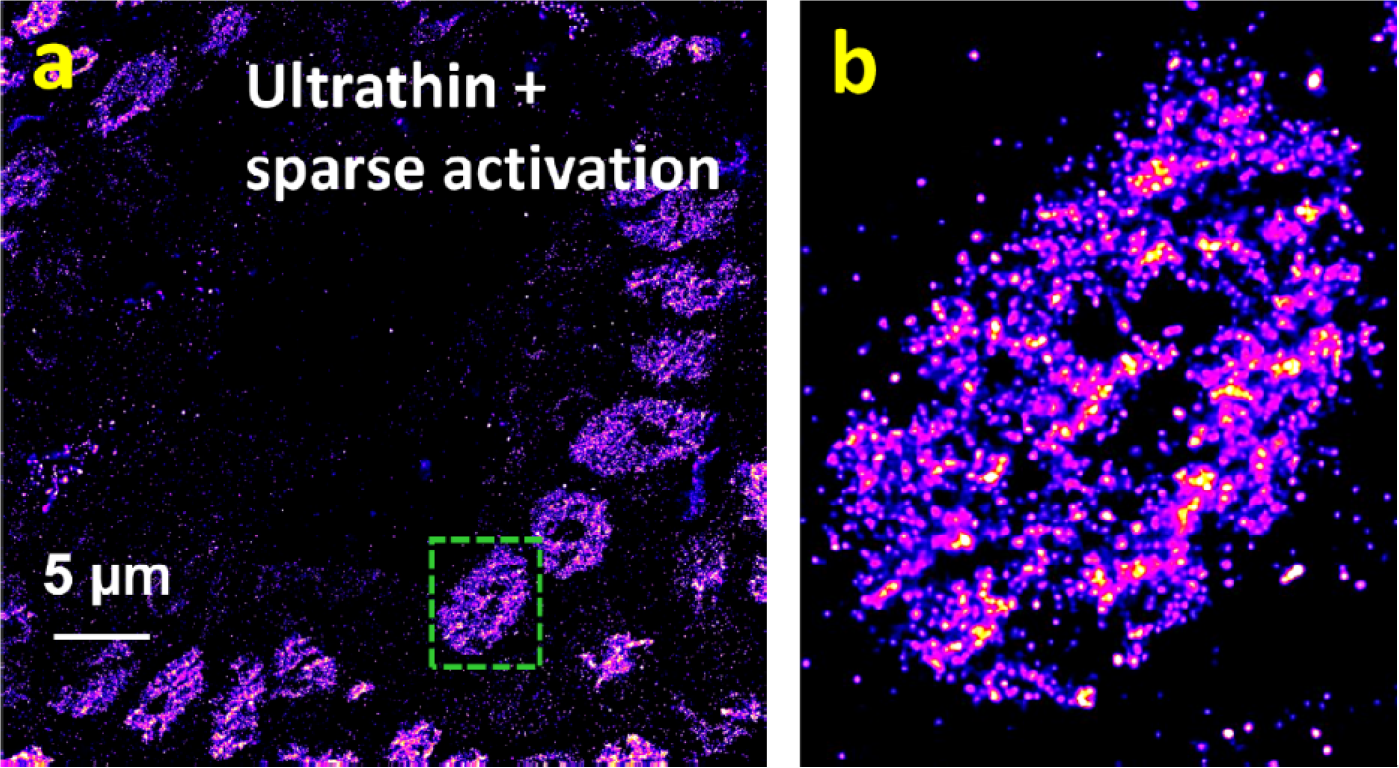
(**a**) The reconstructed super-resolution image using experimental dataset of imaging histone mark (H4Ac) labeled with Alexa 647 on an ultrathin frozen intestinal tissue (thickness estimated to be ∼700 nm) sectioned with an ultramicrotome (Reichert Ultracut). The STORM imaging of ultrathin frozen tissue section shows low background (∼200 photons) with sparsely distributed emitters at each image frame. The imaging condition is the same as STORM imaging of cells (described in Methods of main text) with 20 ms acquisition time and accumulated 40,000 frames. The drift correction was performed during the entire imaging process as previously described^1^. (**b**) The zoomed region in the yellow box in (a). The cluster-like structures formed by H4Ac are clearly visible, which resembles the structures in the reconstructed super-resolution image by WindSTORM in Figs. 6(e, e1) in the main text.

**Figure S10.**
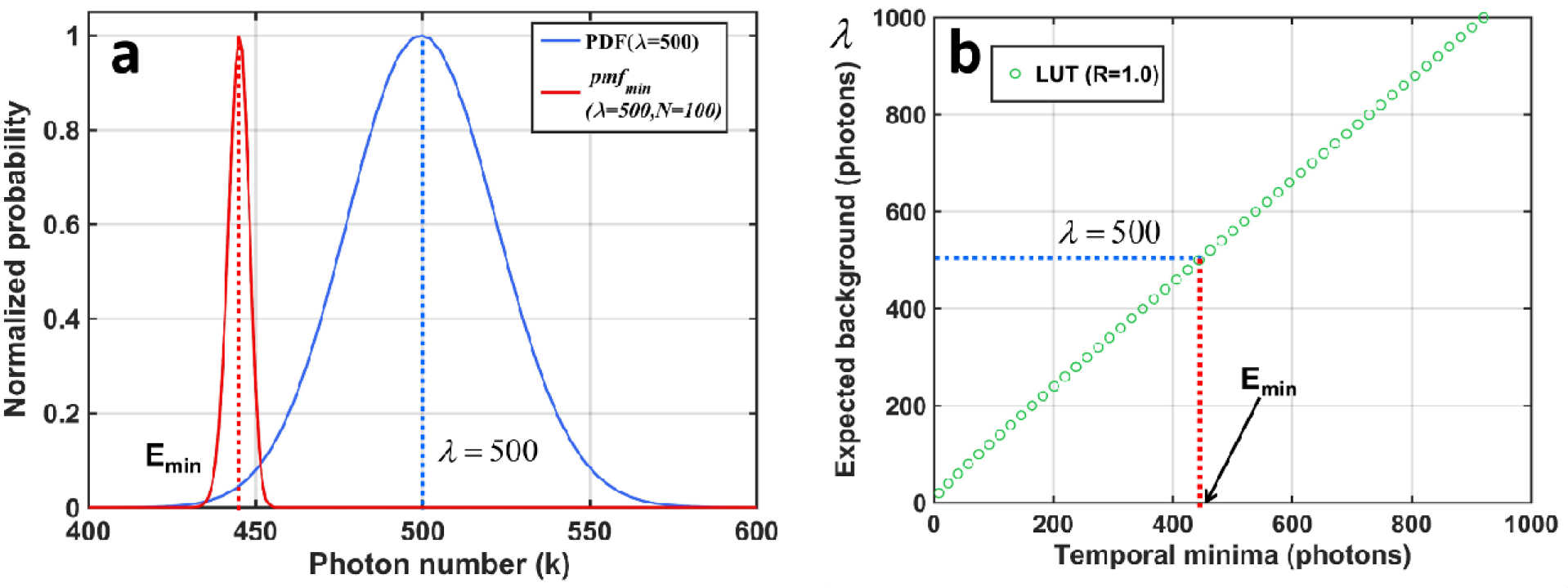
(a) The normalized Poisson distribution function (blue curve) with the expected value (*λ*) of 500 photons and the corresponding normalized probability distribution of the temporal minima (red curve) for N of 100 frames with 3×3 average filter. The dashed line indicates the expected value of the probability distribution of the temporal minima (E_min_). (**b**) An example of the theoretically derived relationship between the temporal minima value to the expected background value based on Eq. S5 when R = 1 (temporally uniform background).

**Figure S11.**
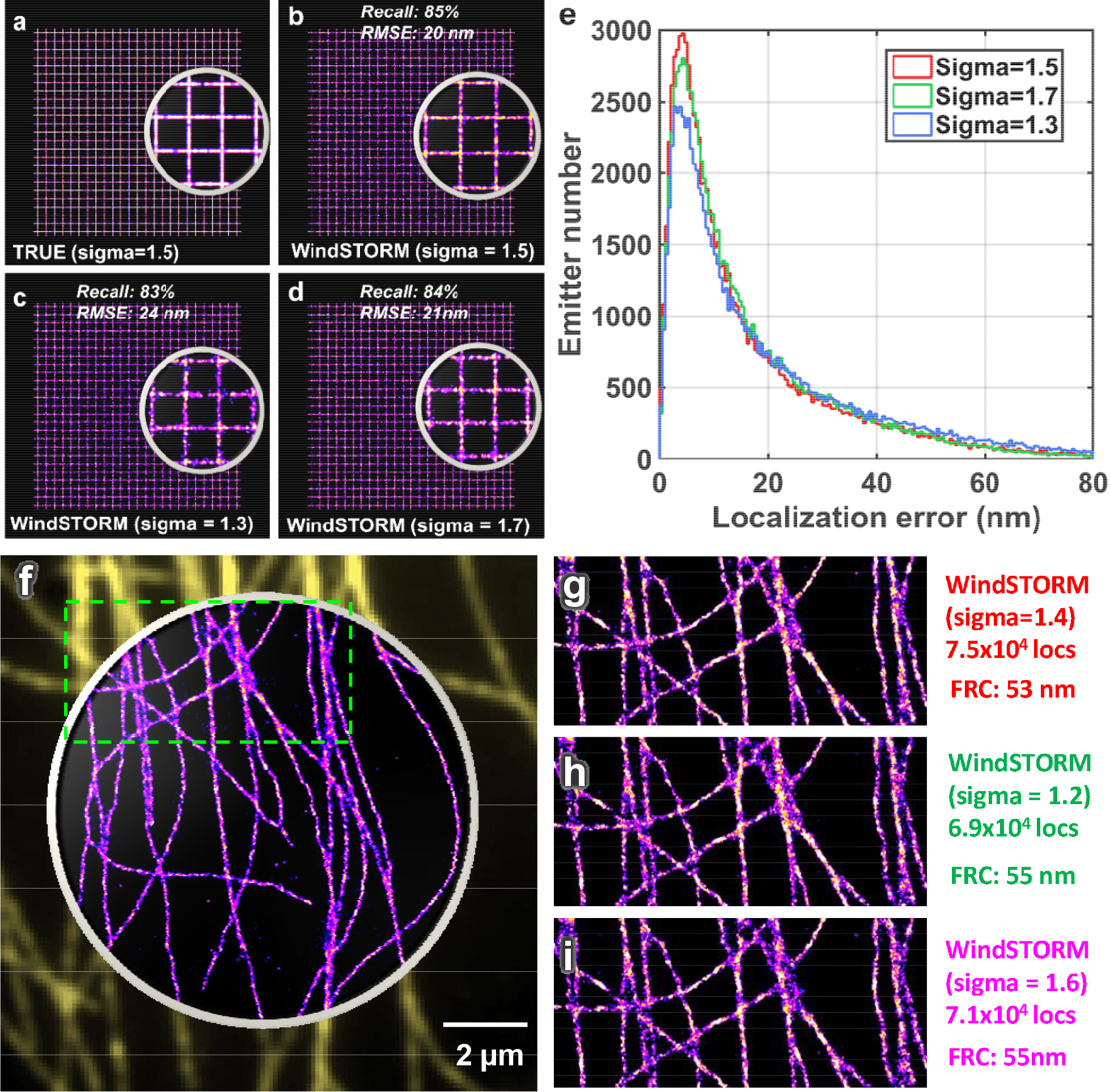
The effect of mismatched PSF width (sigma) between WindSTORM and the actual dataset using both (**a-e**) simulated and (**f-i**) experimental dataset. (**a**) Ground-truth image with PSF width of σ=1.5 pixels; (**b**) The PSF width used in WindSTORM matched with the ground-truth PSF width (σ=1.5 pixels); (**c-d**) mismatched PSF width used in WindSTORM (c) σ=1.3 pixels, and (d) σ=1.7 pixels from the ground truth. (**e**) The histogram distribution of localization errors for matched (red) and mismatched PSF width in WindSTORM (green and blue) with the ground-truth PSF width. The mismatched PSF width results in reduced localization precision by 3-4 nm and reduced emitter recall rate by 1-2%. (**f**) The experimental data of wide-field and super-resolved image reconstructed by WindSTORM. The measured PSF width of the experimental dataset is 1.4 pixels. (**g-i**) The reconstructed super-resolution images by WindSTORM with (g) matched PSF width (σ=1.4 pixels) and (**h-i**) mismatched PSF width (σ=1.2 and σ=1.6 pixels). The FRC resolution is reduced by 2 nm due to the mismatched PSF width.

